# Multimodal tubulin binding by the yeast kinesin-8, Kip3, underlies its motility and depolymerization

**DOI:** 10.1101/2021.10.12.464151

**Authors:** Hugo Arellano-Santoyo, Rogelio A. Hernandez-Lopez, Emma Stokasimov, Ray Yu-Ruei Wang, David Pellman, Andres E. Leschziner

**Author notes:** Equal contribution (Hugo Arellano-Santoyo, Rogelio A. Hernandez-Lopez).

## Abstract

The microtubule (MT) cytoskeleton is central to cellular processes including axonal growth, intracellular transport, and cell division, all of which rely on precise spatiotemporal control of MT organization. Kinesin-8s play a key role in regulating MT length by combining highly processive directional motility with MT-end disassembly. However, how kinesin-8 switches between these two apparently opposing activities remains unclear. Here, we define the structural features underlying this molecular switch through cryo-EM analysis of the yeast kinesin-8, Kip3 bound to MTs, and molecular dynamics simulations to approximate the complex of Kip3 with the curved tubulin state found at the MT plus-end. By integrating biochemical and single-molecule biophysical assays, we identified specific intra- and intermolecular interactions that modulate processive motility and MT disassembly. Our findings suggest that Kip3 undergoes conformational changes in response to tubulin curvature that underlie its unique ability to interact differently with the MT lattice than with the MT-end.

## Introduction

The kinesin family of microtubule (MT)-based motors perform a variety of cellular functions, including organelle transport, cell division and cilia formation. This functional diversity results from the combination of a conserved core motor domain and ATP-dependent catalytic cycle with family-specific domains adapted for specialized functions. One important role of kinesins is in the regulation of MT dynamics and MT length, as underscored by the contribution of the kinesin-8 and kinesin-13 families to accurate chromosome segregation, for example (Bakhoum et al., 2009, Stumpff et al., 2008, Zhang et al., 2019).

The role of kinesin-8s in regulating the MT length of structures like the mitotic spindle and cilia is conserved across eukaryotes, (Gupta et al., 2006, Niwa et al., 2012, Savoian and Glover, 2010, Stumpff et al., 2008, Su et al., 2011, Varga et al., 2009, Du et al., 2010, West, 2001). Unlike other families, Kinesin-8s can move along MTs and reside for prolonged periods at the MT plus-end, where they accumulate and promote MT depolymerization (Grissom et al., 2009, Niwa et al., 2012, Varga et al., 2009, Su et al., 2011), (**Figure 1A**). Therefore the plus-end concentration and depolymerase activity of kinesin-8 becomes proportional to MT length (Su et al., 2011, Gupta et al., 2006, Varga et al., 2009, Arellano-Santoyo et al., 2017).

**Figure 1.**
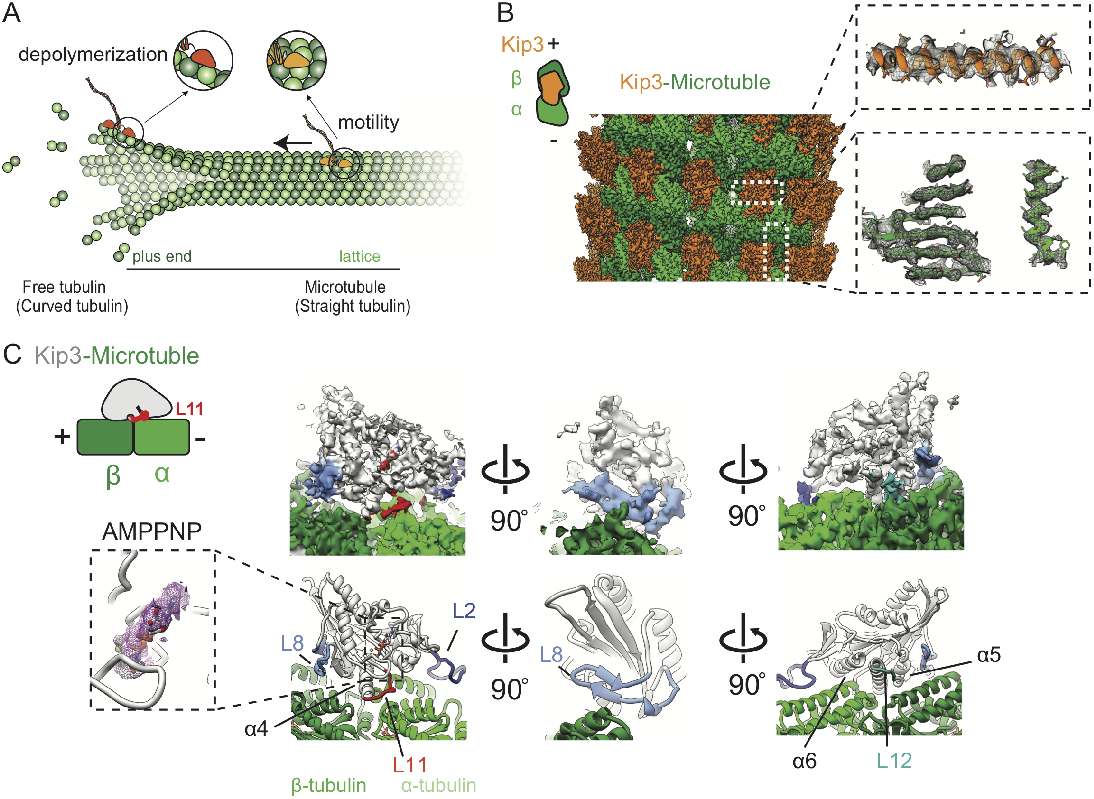
Cryo-EM reconstruction and molecular model of the yeast Kinesin-8 Kip3 (AMPPNP) bound to Taxol-stabilized microtubules. **(A)** Cartoon of Kip3’s activities. On the microtubule (MT) lattice Kip3 (in light orange) is a processive plus-end directed motor. At the plus-end Kip3 (in dark orange) binds tightly to curved tubulin dimers leading to MT depolymerization. **(B)** Three-dimensional reconstruction of monomeric Kip3_438_ bound to Taxol-stabilized MTs. The map has been sharpened and filtered to the calculated overall resolution of 3.9 Å (See **Figure S1**). α- and β-tubulin are shown in light and dark green, respectively. Kip3 is shown in light orange. The MT plus (+) and minus (−) ends are indicated. Inset: Representative densities of α helices and β sheets in the tubulin (in green) and Kip3 (in light orange) regions. **(C)**Segmented density (top) of Kip3 bound to an αβ tubulin heterodimer in Taxol-stabilized MTs and corresponding atomic model (bottom). Loop 11 (L11) is shown in red, loop 12 (L12) is in green and loop 8 (L8) is in light blue and loop2 (L2) in dark blue. The inset shows density (purple mesh) in the nucleotide binding site of Kip3 that fits an AMPPNP molecule.

To regulate MT length, kinesin-8s switch from motility on the lattice to depolymerization at the MT plus-end. This switch in activity arises from preferential binding to different conformations of tubulin; yeast (Kip3) and human (Kif19A) kinesin-8s have been shown to bind more tightly to the curved tubulin state found at the plus-end than to the straight tubulin state located within the MT lattice (Arellano-Santoyo et al., 2017). Several structural elements of the kinesin-8 motor domain have been implicated in mediating MT depolymerization activity. The Loop 2 (L2) and Loop 8 (L8) from human kinesin-8 (Kif18A and Kif19A) extensively contact the MT and have been proposed to control plus-end depolymerization (Peters et al., 2010, Locke et al., 2017, Wang et al., 2016a). However, although the L2 of Kip3 contributes to MT binding and motor processivity, it was found to be dispensable for depolymerization (Arellano-Santoyo et al., 2017). Instead, Loop 11 (L11) plays a critical role in both preferential binding to curved tubulin and depolymerase activity (Arellano-Santoyo et al., 2017). However, how L11 and other regions of the kinesin-8 motor interact with tubulin to mediate the switch from motility to disassembly of the MT plus-end remains poorly understood.

Here, we use cryo-electron microscopy (cryo-EM), molecular dynamics simulations, single-molecule imaging, and mutational analysis of yeast kinesin-8/Kip3 to investigate the mechanism of its interaction with two conformational states of tubulin, straight and curved. We obtained a cryo-EM structure of Kip3 bound to MTs at 3.9 Å, allowing us to identify residues in L8 and L11 that contribute to long-range motility on MTs and to depolymerization activity. Additionally, our cryo-EM structure analysis suggests that despite the presence of an ATP analog (AMPPNP) and the canonical local interactions responsible for nucleotide binding, Kip3 adopts a state similar to the standard kinesin-1 APO conformation when bound to MTs. In contrast, molecular dynamics simulations suggest that when bound to curved tubulin, Kip3 adopts a conformation akin to the kinesin-1 ATP conformation. Finally, functional assays with ATP analogues suggest that MT-depolymerization by Kip3 occurs in a post-hydrolysis ATP state (ADP-Pi). Thus, differential contacts between Kip3 and the MT lattice and the plus-end result in distinct conformations of the entire motor domain that differ from kinesins specialized in motility, such as the kinesin-1 family. We propose a model wherein kinesin-8 undergoes conformational changes in response to tubulin curvature, triggering the switch between motility and depolymerization.

## Results

### Cryo-EM structure of Kip3 bound to MTs

Previous structures of kinesin-8s have given insights into unique adaptations of this family of motors but have been limited by resolutions of ~6-7 Å, precluding visualization of residue-level interactions of kinesin-8s with MTs (Locke et al., 2017, Wang et al., 2016a), Kinesin-8s have large flexible domains and have been difficult to purify under conditions compatible with structural characterization. We overcame these limitations by using a minimal monomeric Kip3 construct (Kip3_438_), which recapitulates the dose-dependent MT depolymerization activity of full-length Kip3 (Arellano-Santoyo et al., 2017). We obtained first a cryo-EM structure of Kip3_438_ bound to GDP-Taxol-stabilized MTs in the presence of the slowly hydrolysable ATP analog adenylyl-imidodiphosphate (AMPPNP), at an overall resolution of 3.9 Å (**Figure 1B,C, Figure S1A and Table S1**).

The majority of the Kip3_438_ sequence is resolved in the structure, although some regions distal to the MT have lower resolutions (**Figure S1B**). However, we were able to identify prominent features in the density, such as the Taxol binding pocket (**Figure S1C)**. We used Rosetta to build atomic models into the density map **(Figure 1C, Figure S1D-F and Table S1**) and used these models for subsequent analysis. The cryo-EM density shows that Kip3 contacts the MT at the αβ-tubulin intradimer interface, primarily through its α4, α5 and α6 helices, similar to other kinesins (**Figure 1C, Figure S1E**) (Sindelar and Downing, 2010, Goulet et al., 2012, Gigant et al., 2013, Shang et al., 2014, Locke et al., 2017, Atherton J, 2014). Two additional prominent contacts with the MT involve Kip3’s L2, as shown previously (Locke et al., 2017, Wang et al., 2016a), and L8 (**Figure 1C**). The nature of the interactions mediated by these structural regions will be discussed in the coming sections.

Kip3 exhibits different activities on the MT lattice and the plus-end. A major difference between these MT structures is that at plus-ends, tubulin is bound to GTP. This gives rise to a region termed the “GTP-cap” that differs from the GDP-bound tubulin in the lattice. To study the interactions of Kip3 with GTP-bound MTs, we used MTs stabilized by the slowly hydrolysable GTP analog, GMPCPP, which mimics the GTP-like state of tubulin (Alushin et al., 2014). GMPCPP-stabilized MTs, unlike GDP-bound, Taxol-stabilized MTs, are a substrate for Kip3-mediated depolymerization (Arellano-Santoyo et al., 2017, Gupta et al., 2006, Varga et al., 2006).

We solved a cryo-EM structure of Kip3 bound to GMPCPP-stabilized MTs, in the presence of AMPPNP, to an overall resolution of 4.1 Å (**Figure 2A, Figure S2 and Table S1**). Overall, the motor backbone conformation of Kip3 was similar to that on GDP-Taxol MTs (RMSD 0.72 Å) (**Figure 2B,C and Figure S2A-F**), however we found some differences in specific domains that mediate interactions between Kip3_438_ and the MT lattice. These domains are L8 and L11 and differed by root mean squared deviations (RMSD) between 1.5 Å and 3 Å (**Figure 2C and Figure S2E**). Upon model building, different sets of interactions between L8, L11 and tubulin residues where observed (**Table S2, Figure S2G-J**), however higher resolution studies will be required to further investigate the relevance of these differences.

**Figure 2.**
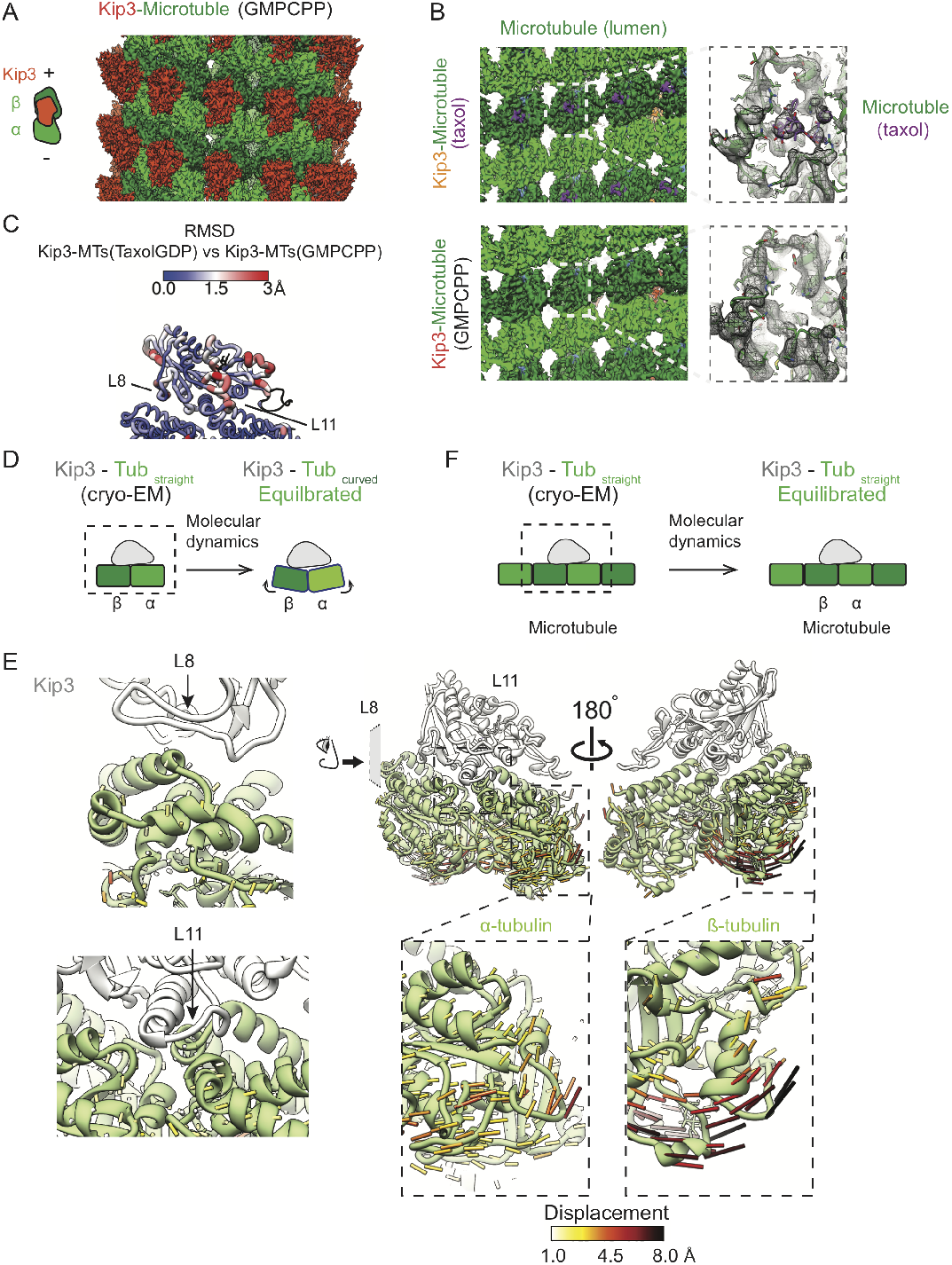
Cryo-EM reconstruction of Kip3 (AMPPNP) bound to GMPCPP-stabilized MTs and model of Kip3 bound to curved tubulin dimers from targeted molecular dynamics simulations. **(A)** Three-dimensional reconstruction of monomeric Kip3_438_ bound to GMPCPP-stabilized MTs. The map has been sharpened and filtered to the calculated overall resolution of 4.1 Å (See **Figure S2**). α- and β-tubulin are shown in light and dark green, respectively. Kip3 is shown in dark orange. The MT plus (+) and minus (−) ends are indicated. **(B)** Cryo-EM maps of Kip3 (AMPPNP) bound to either GDP-Taxol- or GMPCPP-stabilized MTs viewed from the MT-lumen. The Taxol binding site in β-tubulin has been highlighted in both reconstructions and close-up views of the highlighted areas are shown in the insets. The Taxol density is shown in purple. **(C)** Cα atom RMSD map between the top models for Kip3-MT(GDP-Taxol) and Kip3-MT(GMPCPP). L8 and L11, at the Kip3-tubulin interface show the largest RMSD (See **Figure S3**). **(D)** Schematics for approximating the Kip3 - curved tubulin state using targeted molecular dynamics (TMD). Cartoon of TMD simulation used to model changes in Kip3 binding to tubulin from its straight form (left) to its curved form (right). We used our MT (GDP-Taxol) cryo-EM reconstruction as the source for the straight tubulin conformation. To approximate the structure of Kip3 on curved tubulin, the straight conformation of tubulin was used as a starting point for TMD simulations and the curved tubulin Cα positions were obtained from the structure of tubulin in complex with Kinesin-1 and DARPIN (PDB 4HNA). **(E)** Schematics of control MD simulations. MD simulations were performed maintaining the straight tubulin conformation during the entire trajectory. A 4×3 array of tubulin monomers (corresponding to 3 protofilaments with 2 dimers each) was simulated to account for longitudinal and lateral interactions; only one MT-protofilament is shown for clarity. **(F)** Close up views of the Kip3-tubulin complex showing the displacement vectors for the transition from straight to curved tubulin during TMD simulations, colored according to their magnitude (in Angstroms). The insets show expanded views of the indicated regions.

### Molecular Dynamics simulations suggest that Kip3 makes more contacts with the curved form of tubulin

At MT-ends, tubulin subunits lack a full set of stabilizing lateral interactions found in the lattice, and thus adopt a curved conformation similar to tubulin in solution, with a curvature of ~12° compared to straight tubulin found in the lattice (Brouhard and Rice, 2014). Thus, a molecular understanding of Kip3’s plus-end depolymerization mechanism requires a comparison of Kip3 bound to the MT lattice (tub_straight_) versus curved tubulin (tub_curved_, free tubulin).

To characterize the interface between Kip3 and tub_curved_ we attempted to obtain a crystal structure of the Kip3-tubulin complex but were unsuccessful. Therefore, we instead generated a model of Kip3 bound to tub_curved_ using Targeted Molecular Dynamics (TMD) (Schlitter et al., 1994, Phillips et al., 2005). In TMD, a subset of atoms, in this case the Cα atoms of the tubulin heterodimer, is guided towards a target conformation by using steering forces, while the full atomic interactions are computed via molecular dynamics force fields (Schlitter et al., 1994, Phillips et al., 2005). This approach has been useful to generate hypotheses about protein-ligand unbinding kinetics, allosteric mechanisms of transport, and conformational rearrangements during folding or unfolding processes (Schlitter et al., 1994, Provasi et al., 2009, Cheng et al., 2006, van der Vaart and Karplus, 2005). Using TMD, we obtained a model for the Kip3-tub_curved_ complex that allowed us to generate hypotheses that we tested with biochemical and functional assays.

The binding interface between curved tubulin and kinesin-8 was modeled by TMD; we started from our cryo-EM structure of Kip3 bound to MTs and used existing X-ray crystal structures of kinesin-1 bound to curved tubulin as the target. Specifically, we used the straight tubulin conformation from our GDP-Taxol-stabilized cryo-EM structure (tub_straight_) as the starting point since the Kip3-tubulin backbones are very similar in our two cryo-EM structures and the Taxol-stabilized structure was of higher resolution. The curved tubulin conformation (tub_curved_) from the kinesin 1-tubulin-DARPIN complex (PDB 4HNA) (Gigant et al., 2013) was used as the end-point for the TMD trajectory (**Figure 2D**). Importantly, in our TMD simulations the steering forces were only applied to the Cα atoms of tubulin. Thus, Kip3 atoms were not steered by any additional force field during these simulations and any changes observed in Kip3 should be driven by the tubulin conformational changes resulting from the TMD force field (**Figure 2E)**. Because force fields for ATP analogs have not been standardized, we carried out the simulations with ATP in the Kip3 binding pocket. As a control, we performed TMD simulations with tub_straight_ as the start and end-point (**Figure 2F**). We then analyzed the changes in the molecular interactions between ATP-bound Kip3 and tubulin in the tub_straight_ or tub_curved_ conformations.

Our TMD simulations show different interactions between Kip3 and tubulin in the curved and straight conformations. Overall, Kip3 appears to form more interactions with tub_curved_ than tub_straight_: 206 vs. 181 unique salt bridges and 23 vs. 18 unique hydrogen bonds that lasted more than 10% of the 30 ns of simulated time (termed occupancy) (**Tables S3-S5**). Of particular interest among the different interactions were those involving Kip3 L8 (**Figure 3A,B**), which formed twice as many long-lived interactions with β-tubulin (see Methods) in tub_curved_ relative to tub_straight_ (10 vs. 5 hydrogen bonds). We also observed that Kip3 L8 adopted different conformations relative to β-tubulin’s helix 12 (H12) in tub_curved_ and tub_straight_: the center of mass of Kip3 L8 is ~3 Å closer to β-tubulin’s H12 in the curved conformation (**Figure 3C**). Concordantly, an increase in the distance between L8 and the core β-sheet of Kip3, measured by the distance between the beta carbons of F333 and H268, is also observed (**Figure 3D)**. Additionally, we observed several different interactions between positively charged residues in L11 and negatively charged residues in straight and curved tubulin (**Figure 3A,E,F**), which we discuss below. Our simulations suggest that Kip3’s recognition of curved tubulin at MT-ends is primarily mediated by residues in L8 and L11, consistent with Kip3 having a 4.5-fold higher affinity for soluble tubulin than for the MT lattice (Arellano-Santoyo et al., 2017).

**Figure 3.**
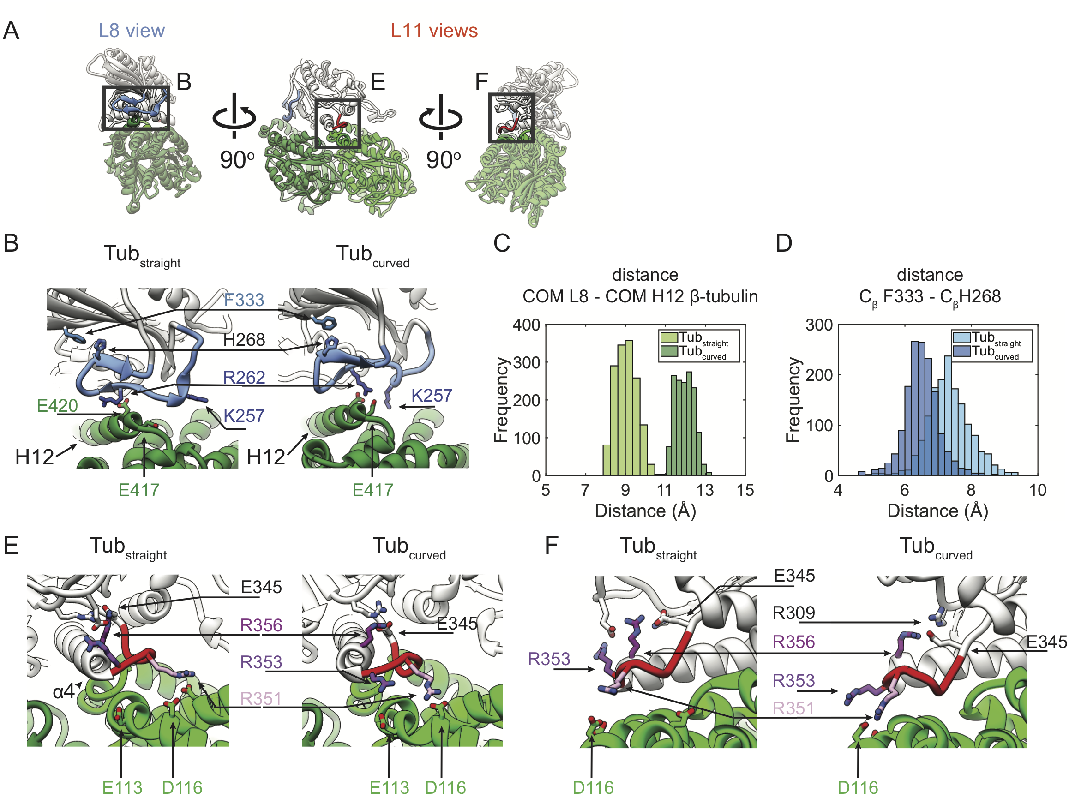
Molecular dynamics simulations identify different interactions between Kip3 and either straight or curved tubulin. **(A)** Three views of the Kip3-tubulincurved model obtained from MD simulations. The boxes show the areas displayed in B, E and F. **(B)** Kip3 L8, colored in light blue, binds differently to curved and straight tubulin. Close-up views of Kip3’s L8 on straight (i.e. MT) (left) and curved (right) tubulin. The residues F333, H268, R262 and K257 are shown in stick representation. **(C)** Kip3 L8 is further away from tubulin in the curved tubulin state. Histogram of the distance distribution between the center of mass (COM) of L8 and the COM of H12 in β-tubulin. **(D)** F333 and H268 are closer together on curved tubulin. Histogram of the distance distribution between the beta carbons (C_β_) of F333 and H268 during the 30 ns MD trajectories of Kip3 on straight (MT) and curved tubulin. **(E,F)** Kip3 L11 residues bind differently to straight and curved tubulin in TMD simulations. Views of the Kip3’s loop 11 bound to straight (left) and curved (right) tubulin as observed in our TMD simulations. Key residues involved in interactions are shown as sticks and color coded in shades of purple.

### Interactions involving loops 8 and 11 play key roles in Kip3’s velocity and long-range processivity

The cryo-EM structures of the Kip3-MT complex (**Figure S3G-J, Table S2**) together with our simulations (**Figure 3A, Tables S3-S5**) suggested three sets of inter- and intra-molecular interactions that might underlie Kip3’s processive motility.

First, there are intermolecular electrostatic interactions made by residues in L8 with residues in helix 12 (H12) in β-tubulin (**Figure 4A**), and by residues in L11 with residues in α-tubulin (**Figure 4B, C**). Mutating residues in L8 (K257A, R262A) or L11 (R351A, R353A) involved in these electrostatic interactions decreased motor processivity (**Figure 4D,F**) and increased velocity (**Figure 4E,G**). These results highlight the role of positively charged residues in modulating kinesin motility on the MT lattice, similar to what has been observed in other kinesins (Fourniol and Moores, 2010).

**Figure 4.**
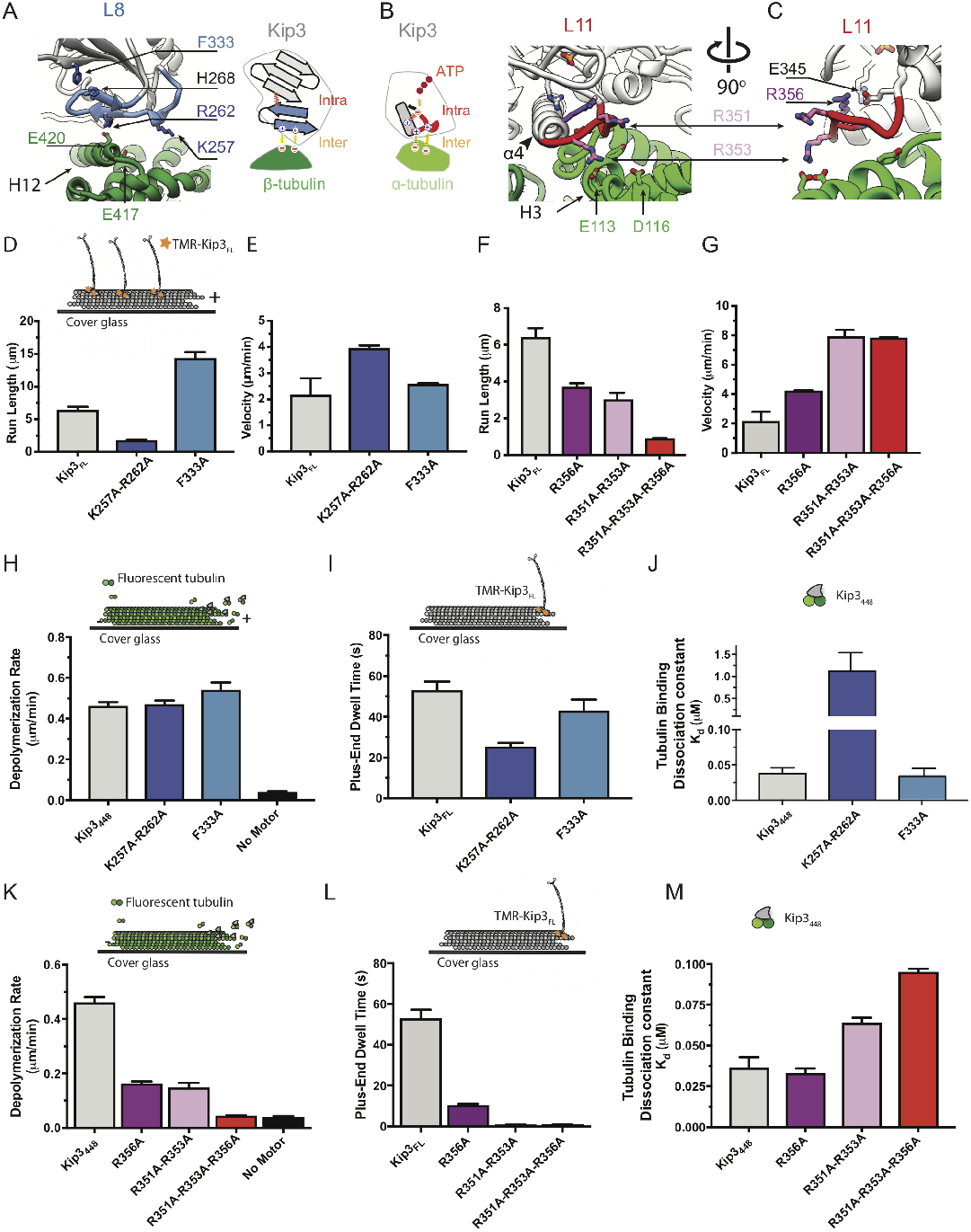
Roles of residues in Kip3’s L8 and L11 in motility and depolymerization. **(A)** View of residues in Kip3 L8 interacting with residues in β-tubulin. The residues F333, H268, R262 and K257 in Kip3 are shown in stick representation color-coded in shades of blue to match the histograms in D-E and H-J. A cartoon representation to the right shows the set of inter and intramolecular interactions in which L8 is involved. **(B-C)** Residues in Kip3 L11 interact with α-tubulin. Key residues are shown as sticks and color-coded in shades of purple to match the histograms in F-G and K-M. A cartoon representation to the left shows the set of inter and intramolecular interactions in which L11 is involved. **(D-E)** L8 residues modulate Kip3’s processivity and velocity. Quantification of run length (D) and velocity (E) of Kip3_FL_ (grey), Kip3_FL_^K257A-R262A^ (dark blue) and Kip3_FL_^F333A^ (light blue) (mean +/− SEM). Inset schematically illustrates the experimental design. **(F-G)** Positively charged residues on L11 modulate Kip3’s velocity and processivity. Quantification of run length (F) and velocity (G) of Kip3_FL_ (grey) and Kip3_FL_ loop 11 mutants (mean +/− SEM). L8 residues modulate tubulin binding but not depolymerization. Quantification of depolymerization, plus-end dwell time and tubulin binding of Kip3_FL_ (grey), Kip3_FL_^K257A-R262A^ (dark blue) and Kip3_FL_^F333A^ (light blue) (mean +/− SEM). Insets in panels H-J schematically illustrate the experimental design. **(H)** Depolymerization rates of monomeric Kip3 (Kip3_488_) and Kip3_488_ L8 mutants on GMPCPP stabilized MTs, normalized for MT binding (see methods, mean +/− SEM). **(I-J)** Charge-neutralizing mutations on L8 residues significantly reduce tubulin binding. Tubulin binding constants (K_d_) of Kip3_488_ and Kip3 L8 mutants, measured by biolayer interferometry. L11 arginines are required for MT depolymerase activity and plus-end binding. Insets in panels K-M schematically illustrate the experimental design. **(K)** Depolymerization rates of Kip3_488_ and Kip3_488_ L11 mutants on GMPCPP stabilized MTs (mean +/− SEM). **(L)** Charge-neutralizing mutations of L11 arginines significantly decrease plus-end dwell time (L) and tubulin binding. **(M)** Tubulin Binding constants (Kd) of Kip3_488_ and Kip3_488_ Loop11 mutants (mean +/− SEM).

Second, we noticed an intramolecular interaction between H268 in L8 and F333 in the main β-sheet of Kip3 (**Figure 4A**). We hypothesized that this interaction could constrain the position of L8 and thus the proximity of its residues K257A and R262A to the MT. In support of this idea, disrupting the intramolecular interaction between L8 and the core of Kip3 (via a F333A mutation) increased motor processivity 2-fold (**Figure 4D**) without a significant change in velocity (**Figure 4E**).

Finally, there is an additional intramolecular salt bridge between the kinesin-super family conserved catalytic glutamate E345 and the kineins-8s conserved R356 at the C-terminal end of helix 4 (**Figure 4B,C, Figure S5A**). Based on the structures of other kinesins, this interaction is expected to reduce ATPase activity by limiting the ability of E345 to activate the nucleophilic water molecule for ATP hydrolysis (Parke et al., 2010, Gigant et al., 2013, Cao et al., 2014). We thus predicted that removal of this interaction would enhance MT-stimulated ATP hydrolysis, and consequently increase motor velocity. Indeed, the Kip3 R356A mutant showed ~2-fold higher MT-stimulated ATPase activity (**Figure S4B,C**), a concordant ~2 fold higher velocity (**Figure 4G**), and ~33% decreased processivity on the MT lattice (**Figure 4F**).

Our data show that Kip3 makes several key molecular interactions that increase MT binding and partially autoinhibit the motor’s ATPase cycle. Together, these effects help the motor avoid detachment from MTs, effectively trading motor velocity for increased processivity (**Figure 4F,G**).

### Kip3’s plus-end dwelling and depolymerization require positively charged residues in L11

Our Cryo-EM structures and molecular dynamics simulations suggested that specific residues within the previously characterized Kip3 L11 sequence (Arellano-Santoyo et al., 2017) are responsible for selective recognition of curved tubulin, tight MT plus-end binding, and depolymerization. Consistent with previous observations, the MD simulations showed that residues in Kip3’s L11, particularly R351 and R353, have higher occupancy in their interactions with tub_curved_ when compared to tub_straight_ (**Figure 3E-F, Tables S3-S5**), making them prime candidates for regulating depolymerization activity. Their potential role is supported by additional evidence. First, Kip3 R351 interacts with α-tubulin D116 in our cryo-EM structures (**Figure S3G,H**), and D116 has been shown to be required for Kip3-mediated depolymerization of GTPγS MTs (Arellano-Santoyo et al., 2017). In our simulations, R351’s interaction with α-tubulin D116 showed ~8-fold longer residence time in the tub_curved_ state than in the tub_straight_ state (**Tables S3-S5**). Second, the position corresponding to R353 is conserved as a basic residue (R/K) in kinesin-8 family members (**Figure S5A)**. Mutating both R351 and R353 to alanine reduced the depolymerization rate of Kip3_448_ ~4-fold (**Figure 4K**) and abolished the ability of Kip3 to dwell at the MT plus-end (**Figure 4L**). Kip3_448_^R351A-R353A^ also displayed a 1.8-fold lower affinity for free tubulin, as measured by biolayer interferometry (**Figure 4M**). Therefore, L11 residues, particularly R351-R353, are required for Kip3 recognition of curved tubulin and MT depolymerization.

At the MT-end, Kip3’s L11 additionally contributes to prolonged plus-end dwelling by suppressing the motor ATPase cycle when bound to curved tubulin (Arellano-Santoyo et al., 2017). Therefore, mutations that increase ATPase turnover and Kip3’s velocity on the MT lattice, like R356A (**Figure 4G, Figure S4B,C**), should affect the ability of Kip3 to depolymerize MTs. Indeed, monomeric Kip3_448_^R356A^ showed a ~2.5-fold slower depolymerization rate (**Figure 4K**) and Kip3_FL_^R356A^ had a 5-fold shorter dwell time at the MT plus-end (**Figure 4L**). In contrast, R356 does not appear to contribute to the tubulin-stimulated ATPase activity or binding of the motor to curved tubulin. Kip3_448_^R356A^ showed no difference from wild-type Kip3_448_ in either binding affinity or ATPase activity with free tubulin (**Figure 4M**, **Figure S5B**). Kip3 must transition from ATPase-driven processivity to a strong binding state at the plus-end and L11-R356 is required for this switch.

Our structural analysis suggested additional residues beyond L11 may be involved in Kip3 depolymerization of MTs. We investigated two family conserved residues in L8 (**Figure S5C)**, K257 and R262, that exhibited strong interactions with tubulin and mediate Kip3’s interaction with β-tubulin during motility. Our simulations showed that salt bridges made by Kip3 K257 and R262 with β-tubulin exhibited higher occupancy in curved tubulin (**Tables S3-S5**). As expected, removing these interactions not only affected Kip3’s motility on the lattice (**Figure 4D,E**), but also Kip3’s residence at the MT plus-end and its binding to free-tubulin: a charge-neutralizing mutant (K257A-R262A) showed a ~2-fold reduction in the plus-end dwell time (**Figure 4I**) and ~10-fold weaker affinity (increased K_d_) for free tubulin relative to wild-type Kip3_448_ (**Figure 4J**). Surprisingly, at concentrations resulting in similar levels of Kip3 accumulation at MT-plus ends relative to wild-type Kip3_448_, the K257A-R262A double mutant had no effect on depolymerization rate (**Figure 4H, Figure S4D**), or the free tubulin-stimulated ATPase activity of Kip3 (**Figure S5D**). Thus, while the interaction between L8 and tubulin contributes to both Kip3’s motility and binding to curved tubulin, it appears not to be required for its depolymerization mechanism (see Discussion).

### Kip3’s conformation on MTs is similar to that of APO kinesin-1, despite the presence of AMPPNP in its nucleotide-binding pocket

Previous studies have identified a conserved mechanism where subdomains in kinesin-1, kinesin-3, and kinesin-5 families undergo structural changes during the nucleotide cycle (Cao et al., 2014, Gigant et al., 2013, Goulet et al., 2012, Atherton J, 2014). Two conformations have been described on MTs and tubulin: a “no-nucleotide” (APO) state and a nucleotide-bound (ATP-like) state in the presence of ATP analogs (Shang et al., 2014, Cao et al., 2014, Gigant et al., 2013).

We compared our cryo-EM structures of MT-bound Kip3 to analogous structures reported for kinesin-1 (**Figure 5A**). Surprisingly, despite the presence of AMPPNP in Kip3’s binding pocket (**Figure 1C**), we noticed that the conformation of Kip3’s core (helices α1-α4, α6) largely resembles that of MT-bound kinesin-1 in the APO state (**Figure 5A**). Despite the global similarity to the APO state motor conformation, local interactions, particularly the closure of the conserved structural elements around the nucleotide-binding site typically associated with the ATP state (switch I loop), can be observed in response to AMPPNP. Because we used a minimal Kip3 construct lacking the neck linker, we wondered whether this region might be necessary for these conformational changes associated to nucleotide binding. Therefore, we compared Kip3 and kinesin-1 with the published structures of the kinesin-8, Kif18A, containing the full neck linker (Locke et al., 2017). These structures were solved in both the APO state (in the presence of apyrase) and the ATP-like state (AMPPNP in the ATP-binding pocket).

**Figure 5.**
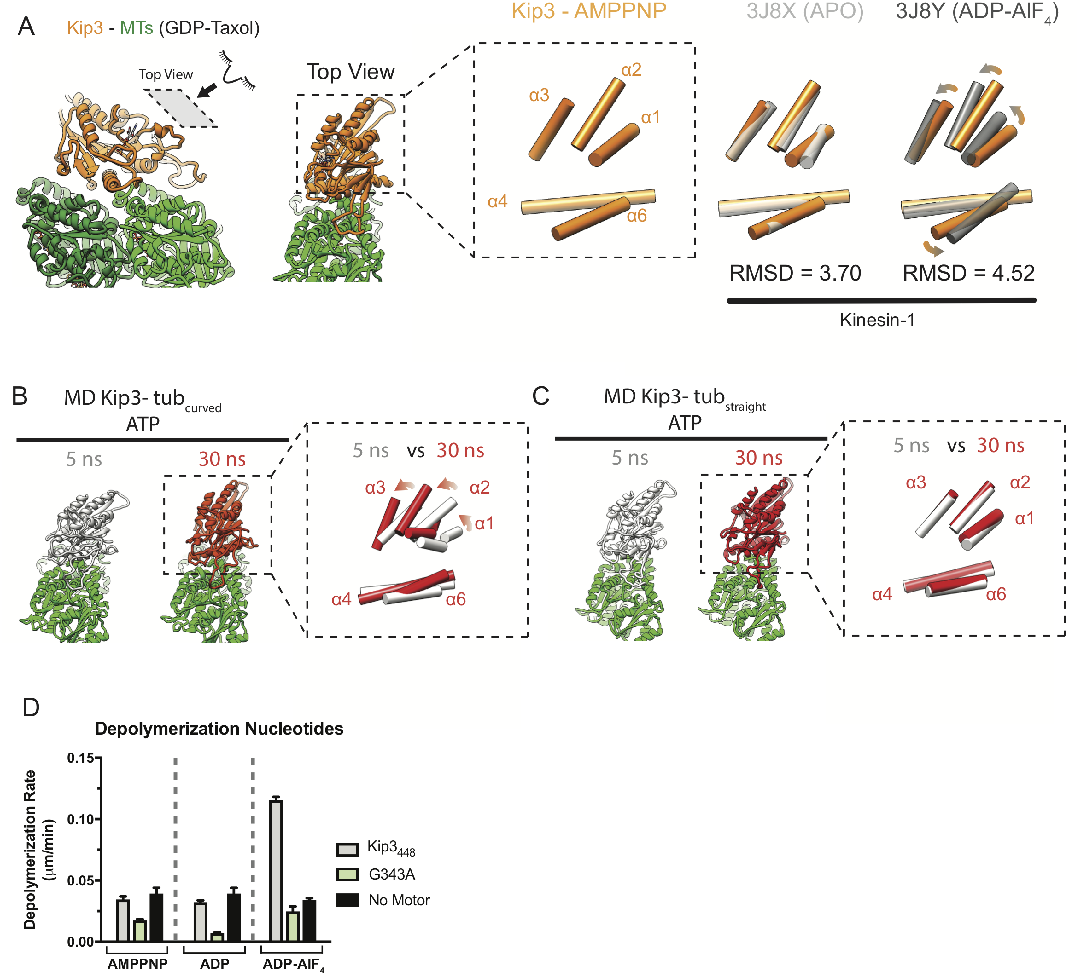
Kip3-AMPPNP adopts an APO-like state on microtubules but switches to an ATP-like state when bound to curved tubulin. **(A)** Kip3-AMPPNP adopts a kinesin-1 APO-like conformation on MTs. Two views of the Kip3-AMPPNP structure on GDP-Taxol (light orange). Inset: Alpha-helices α1-α6 in Kip3 shown from the top view and represented as cylinders. Equivalent views for the superposition of the α1-α6 helices of Kip3 and kinesin-1 APO (white, PDB 3J8X) or kinesin-1 ATP-like (gray, PDB 3J8Y). The atoms in the αβ-tubulin backbone were used for alignment and comparison of the Kip3 and Kinesin-1 structures. **(B)** Molecular dynamics simulations suggest that binding to curved tubulin stabilizes the ATP-like state of Kip3. Snapshots of Kip3 (ATP) bound to tubulin during targeted molecular dynamics simulations (transition from straight to curved tubulin) at 5 ns (white) and 30 ns (dark orange). The inset shows the superposition of helices α1-α6 in Kip3 shown from the top view and represented as cylinders. See **Figure S6B,C** for direct comparison with the kinesin-1 APO and ADP-AlF_4_ structures. **(C)** Binding to straight tubulin is not sufficient for Kip3 to stabilize an ATP-like conformation observed for Kinesin-1. Molecular dynamic simulation snapshots of Kip3 (ATP) bound to straight tubulin (MT) during the simulation trajectory (fixed straight tubulin) at 5ns (white) and 30 ns (dark orange). **(D)** The post hydrolysis state (ADP-AlFl_4_) is Kip3’s depolymerization-competent state. Depolymerization rates of Kip3_488_ (grey) and Kip3_488_^G343A^ (light green) mutant and no motor control (black) on GMPCPP stabilized MTs in the presence of ATP, ADP and ADP-AlF_4_ (mean +/− SEM).

Importantly, and as noted by Locke et al, local conformational changes associated with nucleotide binding such as closure of the binding pocket and neck linker docking can be observed in the Kif18A structures (Locke et al., 2017). However, we find that Kip3’s global conformation (with AMPPNP bound) is similar to both APO and AMPPNP Kif18A structures (**Figure S6A,B**), which are in turn both similar to kinesin-1 in the APO state (**Figure S6C,D**). Thus, our data suggest that in kinesin-8s, local structural rearrangements associated with nucleotide binding can occur in the absence of the larger conformational changes observed in the ATP-like state of kinesin-1.

### Molecular Dynamics simulations suggest that Kip3 adopts the kinesin-1 ATP-like conformation when bound to curved tubulin

Analysis of the targeted molecular dynamics trajectories showed that upon the transition from tub_straight_ to tub_curved_, Kip3’s core switched from the canonical APO-like conformation to ATP-like conformation (**Figure 5B and Figure S7A-C**). In contrast, Kip3’s core conformation remained in the APO-like state in the control simulations on MTs (tub_straight_ -> tub_straight_, **Figure 5C and Figure S7D-F**) even though both simulations had ATP bound in the Kip3 nucleotide pocket. Importantly, the global conformation of Kip3-ATP observed at the end of our control simulations on MTs is similar to our cryo-EM structures of Kip3-AMPPNP on microtubules. These findings suggest that unlike kinesin-1, whose conformation changes are dictated by the nucleotide state of the motor (AMPPNP or ATP), kinesin-8s undergo conformational changes at the motor domain in response to tubulin curvature.

Our molecular dynamics simulations suggest that Kip3 remains in the APO-like conformation when bound to straight tubulin (MTs) despite the presence of ATP and that it undergoes a global conformational change when bound to curved tubulin. Our cryo-EM structures showed that the presence of AMPPNP in the Kip3 nucleotide pocket is not sufficient to support the canonical motor conformational change observed in kinesin-1 in response to microtubule and nucleotide binding. This suggested that these conformational changes may be important for Kip3’s depolymerization activity. Indeed and to our surprise, Kip3 showed no depolymerization activity in the presence of AMPPNP (**Figure 5D**). We considered two possible explanations for this observation: depolymerization by Kip3 either requires a specific conformation that is not fully captured by AMPPNP or lack of activity could be due to tight binding of the monomer onto the MT lattice, thereby reducing the effective soluble Kip3 concentration. However, the assays were conducted at motor concentrations of 250nM (Figure 5D,-figure supplement 3 Kip3_438_) to 1uM (Figure 5D), representing ~2x-9x Kip3: polymerized tubulin (MT) molar excess. We therefore tested various nucleotide state analogs, as well as a Kip3 with a mutation in its switch domain, G343A, rendering the motor unable to undergo conformational transitions in its hydrolysis cycle, regardless of the nucleotide used. We chose to test the well-described mutation G343A, which corresponds to a conserved glycine in the active site of kinesin-1 (G234); this residue forms a hydrogen bond with the gamma phosphate of ATP and triggers a global conformational change upon nucleotide binding (Rice et al., 1999). Thus, the glycine-to-alanine mutation traps kinesin-1 in a pre-hydrolysis conformation, structurally similar to the APO state (Rice et al., 1999). We tested depolymerization by Kip3_448_ or Kip3_448_^G343A^ in the presence of AMPPNP, ADP, or ADP-AlF_4_. For Kip3_448_ neither AMPPNP nor ADP were able to promote depolymerization, but ADP-AlF_4_ supported Kip3 depolymerization activity (**Figure 5D**). Kip3_448_^G343A^ was unable to depolymerize GMPCPP-stabilized MTs in all nucleotide states tested, including ADP-AlF_4_ (**Figure 5D**). Given that ATP and ADP-AlF_4_ support depolymerization in Kip3_448_, but neither AMPPNP nor ADP do (**Figure 5D and Figure S8**), suggests that the depolymerization-competent state of Kip3 is a post-hydrolysis state but prior to phosphate release. Taken altogether, our findings indicate that Kip3-mediated microtubule depolymerization occurs in the ADP.P_i_-like state, which is stabilized upon binding to curved tubulin at the MT plus-end.

## Discussion

Here, we present a structural and functional analysis of the motility and MT depolymerization activities of the yeast kinesin-8, Kip3. We solved two cryo-EM structures of Kip3 bound MTs—Taxol- and GMPCPP-stabilized— and used Molecular Dynamics simulations to model Kip3 bound to curved (free) tubulin and compare it with our cryo-EM structures in the microtubule-bound state. Our structural analysis revealed differences in the interaction of Kip3 with straight or curved tubulin, largely mediated by L11 and L8, that underlie its ability to walk on (**Figure 6B**) and depolymerize MTs (**Figure 6C**). Furthermore, our structural analysis showed that Kip3’s adopts a global conformation on MTs that resembles the APO-like conformation of kinesin-1, despite the presence of an ATP analog in the motor’s nucleotide binding pocket. We propose that conformational changes in Kip3 in response to tubulin curvature enable it to switch between the seemingly opposing functions of processivity and MT depolymerization.

**Figure 6.**
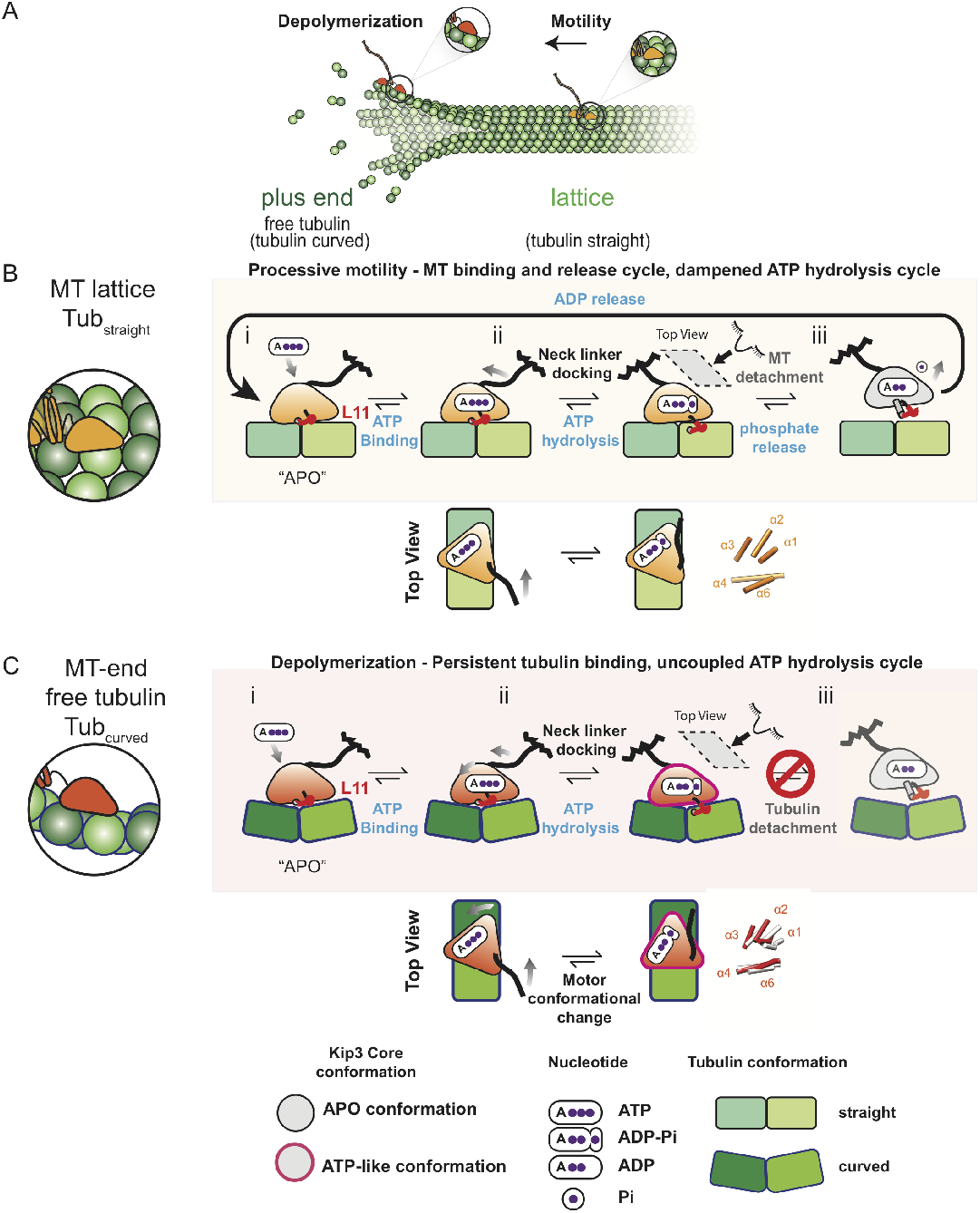
Model for processive motility and depolymerization activity by the kinesin8/Kip3. **(A) The yeast kinesin-8/Kip3 is a multifunctional motor**. Kip3 undergoes processive motility on the microtubule lattice, whereas it strongly binds to the curved tubulin found at microtubule plus-ends, thereby promoting microtubule disassembly. **(B)** On the microtubule lattice, Kip3 undergoes an ATPase cycle similar to other motile kinesins to enable processive motility. After ADP release, (i) ATP binds into Kip3’s nucleotide pocket, promoting strong binding to the microtubule but unlike kinesin-1 the global conformation of the motor remains in a kinesin-1 “APO-like” configuration (i-ii). ATP hydrolysis results in neck linker docking and subsequent phosphate release leads to microtubule detachment (ii-iii). The ATP-like global motor conformation on the microtubule is likely short-lived and is not shown. L11, shown in red, dampens the Kip3 ATPase cycle and kinesin-8 conserved residues in L11, L2 and L8 increase the motor’s affinity for the lattice. **(C)** Kip3’s ATPase cycle is uncoupled from tubulin binding on microtubule ends and free tubulin. (i) Upon encountering curved tubulin at microtubule ends or tubulin in solution, Kip3 L11 adopts a unique conformation. This leads to strong tubulin binding and global structural rearrangements that we propose resemble an ATP-like conformation (as defined for kinesin-1) (i-ii). Changes in L11 binding lead to an inhibited ATPase cycle resulting in a motor tightly bound to tubulin in a post-hydrolysis state (ii-iii), which may be responsible for microtubule depolymerization. The ATP hydrolysis step is highlighted for both microtubule (B, bottom) and tubulin (C, bottom) ATPase cycles. The figure key indicates the motor conformation as defined for kinesin-1, nucleotide state and tubulin conformations.

### L8 is important for motility but not for depolymerization

Structural work on the human kinesin-8 Kif18A and the ciliary kinesin-8 Kif19A have suggested a role for L8 in its interaction with curved tubulin (Peters et al., 2010, Wang et al., 2016a, Locke et al., 2017). Based on a rigid-body fitting of the Kif18A-ADP crystal structure into a 7 Å cryo-EM map of Kif19A-APO bound to MTs (Wang et al., 2016a), it has been proposed that the position of kinesin-8’s L8 adjusts according to tubulin curvature. In this model, L8 retracts towards Kinesin-8’s central ◻-sheet to accommodate binding to curved tubulin. Although an appealing hypothesis, the model was not functionally tested, and the lower resolution of the structures used for the fitting precluded an interpretation in terms of plausible interactions responsible for this L8 retraction. The data we present here show that Kip3’s L8 is involved in interacting with both tub_straight_ and tub_curved_. Disrupting the putative major contacts between L8 (K257A-R262A) and ◻-tubulin reduced Kip3’s binding to both straight and curved tubulin. Thus, our data suggest that the L8-tubulin interaction does not play a major role in discriminating among different tubulin conformations. However, we suggest that an unexpected intramolecular interaction between L8 and Kip3’s core, F333-H268, modulates Kip3’s motility. We predicted that removing it would lead to more frequent or prolongued interaction of L8 residues with residues in H12 of ◻-tubulin (**Figure 4A**). Indeed, disrupting this interaction affected Kip3’s motility on MTs (**Figure 4D**), but not depolymerization (**Figure 4H**). Therefore, our structural and functional data support a role for L8 in motility and tubulin binding but not in depolymerization.

### Kip3 undergoes a global conformational change in response to tubulin curvature

Cryo-EM and X-ray crystallography structures have shown that kinesin-1 adopts two major global conformations during its mechanochemical cycle on both MTs and free tubulin: APO and ATP-like. These conformations differ by local changes in the nucleotide-binding pocket, the neck linker position and a global rotation of kinesin’s upper domain (Shang et al., 2014, Cao et al., 2014, Gigant et al., 2013). Cryo-EM structures at ~6-7 Å resolution of the human kinesin-8, Kif18A, on GDP-Taxol-stabilized MTs and in the presence of apyrase and AMPPNP, showed that kinesin-8s undergo local conformational changes around the nucleotide binding site similar to those observed in kinesin-1 (Locke et al., 2017). The presence of AMPPNP leads to local neck linker docking and closure of the nucleotide binding pocket (Locke et al., 2017). However, in contrast to kinesin-1, Kip3 and Kif18A did not show the characteristic global conformational change associated with the movement of the upper domains. Instead, Kip3 adopts a state that closely resembles the APO-like conformation (**Figure 5A, 6C-middle**) despite the presence of AMPPNP in the nucleotide-binding pocket (**Figure 1C**). Importantly, the similarity of the global conformation between the Kif18A-AMPPNP structure and ours (**Figure S6A**) suggests that the observed APO-like global conformation was not due to the absence of a neck linker in the Kip3_438_ construct but rather due to a conserved kinesin-8 conformation in the MT-bound state.

Although Kip3’s global conformation on MTs closely resembles that of kinesin-1’s APO-like conformation, MD simulations suggest that Kip3 adopts an ATP-like conformation when bound to curved tubulin (**Figure 5B, 6C**). Importantly, our MD simulations did not impose any steering force on Kip3; its conformational changes were a consequence of the changes in the tubulin conformation in its trajectory towards the curved state. Additionally, Kip3 was only able to depolymerize MTs in the presence of ATP and ADP-AlF_4_ (**Figure 5D**). The G343A mutant, which is thought to trap the motor in a pre-hydrolysis state, APO-like conformation, abolished depolymerization activity in all nucleotide states. Furthermore, the E345A mutant, which is thought to trap the ATP-like pre-hydrolysis conformation is capable of depolymerization despite being catalytically inactive (Arellano-Santoyo et al., 2017). Together, these data suggest that depolymerization by Kip3 requires the motor to adopt a unique global conformation, most likely a hydrolysis-like state that resembles the ATP-like conformation of kinesin-1.

### A model for Kip3-mediated MT depolymerization

Previous work has shown that Kip3 regulates MT size by depolymerizing MTs in a length-dependent manner. Kip3 is able to accumulate at MT plus-ends due to its high processivity and tight binding to the curved form of tubulin resulting in plus-end depolymerization (Su et al., 2011, Gupta et al., 2006, Varga et al., 2009, Arellano-Santoyo et al., 2017).

Here, we have shown that Kip3’s high processivity is the result of an increased affinity for tubulin and a dampened MT-stimulated ATPase activity (**Figure 4, Figure S4A,B, Figure 6B**). Our structures suggest that a salt bridge between a basic residue at the C-terminus of 74 (R356) and the catalytic glutamate (E345) might weaken the latter’s ability to activate the nucleophilic water molecule (**Figure 3F**). Although it is unclear if this is a common mechanism of MT-stimulated ATPase activity modulation and thus processivity in kinesin-8s, sequence alignment of the region shows that the basic nature of this residue is highly conserved in members of this family (**Figure S5A**).

Previously, a switch that couples stronger tubulin binding to suppression of ATPase activity in response to tubulin curvature was proposed as the molecular mechanism for Kip3’s ability to depolymerize MTs (Arellano-Santoyo et al., 2017). This stemmed from the observation that a sequence containing the L11 region was responsible for strong tubulin-binding and suppression of tubulin-stimulated ATPase activity. Replacement of this region by the equivalent domain from kinesin-1abolished depolymerization (Arellano-Santoyo et al., 2017). Based on the data presented here, we propose that strong tubulin binding by Kip3 is not sufficient for depolymerization. Instead, both strong tubulin binding and suppression of ATPase activity are needed, and these functions may be uncoupled when bound to free (curved) tubulin (**Figure 6C)**. This is supported by the observation that none of the mutants that showed decreased affinity for soluble tubulin (K257A-R262A, R351A-R353A) (**Figures 4J,M**) showed any changes in the tubulin-stimulated ATPase activity relative to wild-type (**Figure S5A,B**). Furthermore, although the L11-R356A mutation increased the MT-stimulated ATPase activity of Kip3, it did not affect the ATPase activity stimulated by soluble tubulin (**Figure S4B, S5B**).

We propose that an ATP-like (ADP.Pi-like) global conformation of Kip3 bound to curved tubulin, as suggested by the MD simulations, underlies its ability to depolymerize MTs. The importance of the ATP-like conformation in Kip3’s depolymerization is supported by the inability of Kip3_448_^G343A^, a mutant that is locked in the APO-like conformation, to depolymerize MTs (**Figure 5D, 6C**). In addition to a depolymerization competent conformation, a specific nucleotide-state is required for depolymerization as Kip3 was only able to depolymerize MTs in the presence of ADP-AlF_4_ and ATP (**Figure 5D, 6C**). We propose that the low tubulin-stimulated ATPase activity of Kip3 may be due to a prolonged ADP.Pi state; further biochemical assays, however, will be required to test this hypothesis. Taken together, our data suggest that the switch from processive motility to MT depolymerization by Kip3 (**Figure 6A-C**) requires unique contacts between Kip3 and curved tubulin, a conformational switch in the motor domain and a nucleotide-bound, post-hydrolysis state of the motor (**Figure 6B-C**).

Although our structural and functional analyses support this model, we do not yet understand the structural basis for the conformation-dependent uncoupling of Kip3’s ATPase and tubulin-binding. It is unclear why both Kip3 and Kif18A decouple the local conformational changes associated with nucleotide binding from their global conformations. Both Kip3 and Kif18A adopt a global conformation resembling the APO-like state on MTs in the presence of AMPPNP. However, this departure from the conformational changes observed in the canonical catalytic cycle of kinesin-1 is not unique to kinesin-8s. MKLP2, a kinesin-6 involved in cell division, and KLP10A, a depolymerase kinesin-13, show similar global conformations of the motor in the APO and AMPPNP states (Benoit et al., 2018, Atherton J, 2014), as is the case with kinesin-8s when bound to MTs. Whether the conformational transition of Kip3 only occurs on the tubulin-bound state and is the driving force of MT depolymerization will require additional structures of Kip3 bound to curved and straight tubulin in different nucleotide states along with functional assays. Nonetheless, kinesin-8s demonstrate how intermolecular and intramolecular interactionss fine-tune their catalytic cycle and binding properties to result in specialized coexisting functions: highly processive motility along the MT and depolymerization at MT ends, which enabled the emergent property of MT-length regulation.

## Materials and Methods

### Strains and coding sequences for recombinant proteins

Yeast strains were made in the S288C genetic background except those used for protein purification, which were in the W303-1A background. A protease-deficient yeast strain was used for protein expression (Hovland et al., 1989). Expression plasmids for Kip3 purification were previously described (Su et al., 2011, Su et al., 2013). All Kip3 constructs were introduced into a vector with a *GAL1,10* promoter for expression. An N-terminal HALO tag (Promega) for fluorescent labelling was fused to the coding sequence through a GGSGGSLQ linker. Sequence comparisons were performed using ClustalW and CobaltAlignment (NCBI) using the sequences of known kinesin-8s, kinesin-13s and human kinesin-1 as comparison. Point mutations in Kip3 were performed using QuikChange XL (Stratagene). All Kip3_438_ and Kip3_448_ constructs harbor the *Δ*L1 and *Δ*L2 deletions [*kip3 Δ(30-85; 106-134)*]. All constructs were verified by DNA sequencing (Genewiz).

### Kip3 purification protocol

Protein expression was induced by adding 2% galactose for 18h at 30°C prior to harvesting cells and freezing in liquid nitrogen. Cells were disrupted by mechanical force using a blender. The yeast powder was thawed and dissolved in cold lysis buffer (50 mM NaPO_4_, 500 mM NaCl, 10% glycerol, 0.5 mM ATP-Mg, 40 mM imidazole, 2 mM DTT, 1% Triton X-100, protease inhibitor cocktail tablets (Roche), and 2mM PMSF; pH 7.4). Dounce homogenization was used to further lyse cells at 4°C. After centrifugation for 45,000 rpm, 45 min at 4°C, the supernatant was collected and incubated with Ni^2+^ sepharose (GE Healthcare) for 30 min 4°C. The beads were washed 3 times with wash buffer (50 mM NaPO4, 500 mM NaCl, 10% glycerol, 0.1 mM ATP-Mg, 25 mM imidazole, 0.1 mM DTT, protease inhibitor cocktail tablets, and 1 mM PMSF; pH 7.4). The protein was fluorescently labeled with 10uM TMR (dimer constructs) or 30uM TMR (monomer constructs, HALO-TMR; Promega) on beads for 1h at 4°C. Elution off the Ni beads was performed at a rate of <1ml/min with elution buffer (50 mM HEPES, 300 mM NaCl, 50 mM EDTA, 5 mM MgCl_2_, 40mM Imidazole, 50uM ATP-Mg, and 0.1 mM DTT; pH 7.4). The protein eluates were adjusted to 250 mM NaCl and loaded onto a Uno S1 (Bio-rad) ion exchange column, equilibrated with 50 mM HEPES, 250 mM NaCl, and 50 uM ATP-Mg at pH 7.4. The protein was eluted by a linear gradient to 1M NaCl over 10 column volumes. Eluates were supplemented with 10% sucrose and centrifuged at 80,000 rpm for 5 min at 4°C. The supernatant was collected and snap-frozen in liquid nitrogen and stored at −80°C.

### Tubulin polymerization for Cryo-EM

Highly purified, glycerol-free tubulin (Cytoskeleton, Inc.) was resuspended in BRB80buffer (80 mM PIPES-KOH, pH 6.8; 1 mM MgCl2, 1 mM EGTA, 1 mM DTT) to a concentration of 10 mg/mL. To prepare <u>Taxol-stabilized microtubules</u>, tubulin was polymerized with a stepwise addition of Taxol as follows: 20 μL of tubulin stock was thawed quickly and placed on ice. 10 μL of BRB80 supplemented with 3 mM GTP were added and the mixture was transferred to a 37 °C water bath. After 15, 30, and 45 minutes, additions of 0.5, 0.5, and 1.0 μL of 2 mM Taxol were added by gentle swirling. The mixture was then incubated for an additional 1 h at 37°C.

<u>GMPCPP microtubules</u> were polymerized using GMPCPP seeds (20 μM tubulin, 1mM GMPCPP, 1mM MgCl_2_, 1mM DTT in BRB80). A 1 μL aliquot of seeds was transferred to a 37 °C water bath. After 30 minutes, 40 μL of elongation mix (2 μM tubulin, 0.5 mM GMPCPP, 0.5 mM MgCl2, 1 mM DTT in BRB80) were added and the mixture was then incubated for additional 4.5 - 5 hours at 37 °C.

### Grid preparation and imaging

Purified Kip3_438_ protein was buffer exchanged to cryoEM buffer (50 mM Tris-HCl, pH 8.0, 1 mM MgCl2, 1 mM EGTA, 1 mM DTT supplemented with 2mM AMPPNP) and desalted using a ZEBA spin desalting column. The protein was recovered by centrifugation at 15,000 rcf for 2 min. A final spin at 30,000 x g in a TLA 100 rotor (Beckman) for 10 min at 4 °C was carried out to remove big aggregates.

C-flat 1.2/1.3-2C holey carbon grids (Protochips) were glow-discharged for 20 s at 20 mA in a table top plasma cleaner (Electron Microscopy Sciences). Taxol-stabilized MTs were diluted to 0.5 mg/mL in cryoEM buffer supplemented with 100 μM Taxol and GMPCPP microtubules were diluted to 0.25 mg/mL in BRB80 buffer. 4 μL of MTs were added to a grid and allowed to absorb for 30 s. The solution was manually blotted from the side with a torn Whatman #1 filter paper. Next, 4 μL of desalted AMPPNP-kip3 were added and allowed to bind to the MTs for 30 s. The solution was blotted manually again, and the process of addition and blotting of kinesin was repeated for a total of four times. The final blotting was done inside the humidity chamber of a Vitrobot Mark IV (FEI) set at 22° C and 100% humidity. The grids were then rapidly plunged into a liquid nitrogen-cooled ethane slush. Grids were stored in liquid nitrogen until imaging.

Grids were imaged on a FEI Titan Krios (Janelia HHMI Research Campus), equipped with two direct electron detectors, an image corrector for spherical aberration correction, a Gatan Image Filter (GIF), and a high-brightness field emission gun (X-FEG). The two direct electron detectors are a pre-GIF FEI Falcon and a post-GIF Gatan K2 Summit. Images of Kip3 decorating Taxol-MTs or GMPCPP MTs were collected using either a FEI Falcon or a Gatan K2-summit direct detectors, and nominal magnification of 59,000X (Falcon) and 50,000X (K2-summit). Final accumulated electron doses were 25 electrons/Å^2^ for the Falcon camera and 40 electrons/Å^2^ for the K2-camera. Images using the K2-camera were collected in super-resolution mode. The total exposure time was 4 seconds, fractionated into 20 subframes, each with an exposure time of 0.2 s. The calibrated magnifications resulted in images of 1.07 Å/px (Falcon) and 0.52 Å/px (K2 camera in super resolution) on the specimen. Images were recorded using a semi-automated acquisition program, FEI automated software (Falcon Camera) and Serial EM (K2-summit) with a defocus range from 1.5 to 3.5 μm.

Another two data sets of Kip3-decorated GMPCPP Microtubules were collected on a TF30 Polara electron microscope (FEI) operated at 300 kV and equipped with a K2-summit direct detector. Images were recorded in super resolution counting mode at a nominal magnification of 39,000x corresponding to a calibrated super-resolution pixel size of 0.49 Å/px. The total exposure time was 5 s, resulting in a total accumulated dose of 40 electrons/Å^2^ divided into 20 subframes, each with an exposure time of 0.25 s. Images were acquired using Serial EM with a defocus range from 2.0 to 3.5 μm.

### Image processing and three-dimensional reconstruction

Inspection, defocus estimation, microtubule picking, and stack creation were performed within the Appion processing environment (Lander et al., 2009). Images were selected for processing on the basis of high decoration, straight MTs, and the absence of crystalline ice. The contrast transfer function (CTF) was estimated using CTFFIND4 [(Rohou and Grigorieff, 2015) CTFFIND4: Fast and accurate] with a 500 μm step search. A second CTFFIND run with a 100 μm step search was carried out to refine the initial defocus values. Micrographs whose estimated resolution was lower than 8 Å with 0.8 confidence were excluded. Microtubules were manually selected and box files containing square segments of 720 pixels centered around the MT were written at a spacing of 80 Å. Pixel intensities were normalized using XMIPP and segments were decimated 3-fold. Reference free 2D-classification of particles binned by a factor of 3 were subjected to 5 rounds of iterative multivariate statistical analysis (MSA) and multi-reference alignment (MRA) using CAN as described previously (Redwine et al., 2012, Alushin et al., 2014). We also carried out the particle extraction, normalization and reference-free 2D classification using Relion (Scheres, 2012). We found that both methods are reliable to sort MT segments on the basis of degree of decoration and protofilament number (PF) giving roughly the same population distribution of highly decorated 13 or 14 protofilament microtubules (Data not shown). After analyzing the power spectrum and one-dimensional projection of class averages, good quality particles assigned to highly decorated 13 and 14 protofilament MTs were selected for further processing.

For 3D reconstruction we utilized a protocol described previously (Alushin et al., 2014) with some modifications. Synthetic models for a turn of a 13- and a 14-protofilament undecorated microtubules (PDB 1JFF (Lowe et al., 2001)) were used to generate initial low resolution volumes (20 Å) at the calibrated magnification using SPIDER. These volumes were used as references in a multi-model projection matching classification and reconstruction using custom scripts of EMAN2/SPARX (Alushin et al., 2014). Projections along the main microtubule axis are created using an angular step that is decreased incrementally from 4 to 1 degrees. A 10 degree out of plane tilt was also included to create references for alignment.

After each reconstruction cycle, helical parameters for three-dimensional reconstructions were obtained using the hsearch_lorentz program (Egelman, 2000). The helical symmetry was imposed in real space and the protofilament with all the Kip3-tubulin correctly aligned was extracted using a wedge mask. This “correct” protofilament was used to build a new average model with the MT-seam present to start a new refinement cycle.

We included the CTF correction and further sorted the MT segments using a likelihood-based classification recently implemented in FREALIGN (Lyumkis et al., 2013). When the classification converged, the stacks were split into 13 or 14 PF MTs particle stacks for further refinement of the alignment parameters. For refinement in Frealign, the Euler angles were refined first without any symmetry constrains, the helical parameters were determined using the hsearch_lorentz program (Lyumkis et al., 2013) and the symmetry was imposed using a pseudo-helical symmetry operator implemented in FREALIGN without further refinement of helical parameters. In this algorithm, each 80 Å segment is included multiple times in the reconstruction, using the Euler angles and shifts, to generate symmetrically equivalent views. Due to the presence of the seam in the 13 or 14 PF MTs, each segment was inserted 13 or 14 times. An EMAN2 script was used to rebuild the MT containing the correct seam position, as described (Alushin et al., 2014). The best alignment parameters obtained with data binned by a factor of 2 were used as initial parameters to further refine the unbinned data.

The 3D-reconstructions obtained from the data collected with the Falcon camera were used as initial models to process the K2-camera data sets. Dose fractioned super-resolution stacks were aligned to remove beam induced-motion and binned 2×2 using the *motioncorr* UCSF software (Li et al., 2013). After motion correction the sum image was used for initial processing in Appion. The same processing protocol was used as described above. Initially, 3D-reconstructions were obtained using the averaged particles from the 20 subframes of motion corrected images. Particles above the overall average score in Frealign were excluded from the final reconstructions. To accurately estimate the MT seam position we followed a search protocol previously described (Zhang and Nogales, 2015). Briefly, particles within each MT are assigned the consensus euler angles and then an average view of all particles within each MT is generated. For each average, equivalent views considering the multiple protofilaments and alpha-beta registry are created using 40 Å shifts. Each particle and their corresponding Euler angles are tested against the current model and the values with the highest score are assigned as the true seam position. Further refinement using the best particles but averaged from subframes 2-5, 6-10, 10-13 and 14-17 improved the resolution further to the final values. The final resolution of each reconstruction was estimated by calculating the Fourier Shell Correlation of the entire reconstruction or a single tubulin dimer extracted from odd and even volumes as previously described (Alushin et al., 2014).

For the best Kip3-MT Taxol and GMPCPP reconstructions, a local resolution analysis and resolution guided local filtration were performed using ResMap (Kucukelbir et al., 2014) and Blocres (Heymann and Belnap, 2007). The two approaches were consistent with the visual inspection of our reconstruction after sharpening the maps. The resolution of the map containing the Kip3-MT interface is higher than elements away from the MT surface. The average resolution of the 14PF Kip3-MT(Taxol) map calculated from half volumes is 3.9 Å (FSC 0.143 criterion), whereas the tubulin portion is at 3.2 - 4.0 Å, average 3.8 Å (FSC 0.143 criterion) resolution, consistent with our ability to see bulky side chains and the nucleotide densities and the Taxol molecule bound to tubulin. For the Kip3-MT (GMPCPP) the average resolution is 4.1 Å (FSC 0.143 criterion). For visualization of the high resolution features and model building, the final maps were sharpened with a a B-factor of −150 Å^2^ using the program BFACTOR, with the high resolution cutoff determined by the FSC 0.143 criterion.

### Atomic model building and refinement

The general procedure for atomic model building and refinement in cryo-EM density maps using Rosetta were performed as described (Wang et al., 2016b). To prepare initial models for the MT complexes, the atomic coordinates for the tubulin dimers were adopted from the 3.3 Å cryo-EM structure (PDB 3JAT) (Zhang et al., 2015); the ligand coordinates were adopted from other MT structures, including GTP/MG (3J6G), GDP (1JFF), Taxol (3J6G) and GMCPP/MG (3JAT). For Kip3_438_, the starting model was obtained from the Robetta protein structural modeling server (http://robetta.bakerlab.org/). Robetta, which employs the RosettaCM homology modeling pipeline [cite RosettaCM], used PDB 4FRZ as the template and built a full-length model for the Kip3_438_ without using the cryoEM density [cite RosettaCM]. The ligand coordinates AMPPNP/MG taken from 3HQD were then appended to the Robetta Kip3 model.

To capture all inter-domain molecular interactions in the filament, similar to other published MT cryo-EM structures (3JAT and 3J6G), the cryoEM reconstructions of the Taxol-stabilized and GMPCPP-stabilized kip3-MT complexes were segmented to comprise a lattice of 3×3 tubulin units using UCSF Chimera (Pettersen et al., 2004), and were filtered at 3.5 Å and 3.8 Å, respectively. To model/refine the 3×3 lattice of the kip3-MT complexes using Rosetta, symmetry was used to enable energy evaluations of all neighboring interactions around the asymmetric unit (the center of the 3 by 3 lattice).

Multiple rounds of refinement were carried out against one half map (training map), and the other half map (validation map) was used to monitor overfitting based on the procedure described in Wang et al. It is to note that the molecular interactions of ligand-protein were restrained to the initial poses adapted from the high-resolution structures during structure refinement. Finally, we used the half maps to determine a weight for the density map that does not introduce overfitting. Using the weight and with the symmetry imposed, we refined the two kip3-MT complexes using full maps, followed by B-factors refinement. The refined models were validated using MolProbity (Chen et al., 2010) and EMRinger (Barad et al., 2015).

### Molecular dynamics simulations

To gain some insights into differences between the binding of kip3 to straight and curved tubulin we carried out molecular dynamics (MD) simulations. For the straight tubulin state (MT state), a system comprising a kip3 bound to a 3×3 grid of tubulin monomers, derived from our EM reconstruction was embedded in a box of water molecules using the VMD plugin “Solvate” with a 10 Å distance from the closest protein atoms to the box edge. The VMD plugin “Autoionize” was used to randomly place potassium and chloride ions that simulate a final KCl concentration of 0.05 M. All-atom molecular dynamics simulations were performed using the software NAMD 2.9 (Phillips et al., 2005), the CHARMM27 force field with CMAP correction terms (Mackerell et al., 2004) and the TIP3P water model (Jorgensen et al., 1983). Previously derived topology parameters for the GDP, Mg, GTP (Wells and Aksimentiev, 2010) and Taxol (Mitra and Sept, 2008) were utilized during the simulations. The system was minimized for 2,000 steps followed by 30 ns of MD at 300 K. The long-range electrostatic interactions were calculated using the Particle Mesh Ewald method (PME) and the van der Waals interactions were computed with a 10 Å cutoff using periodic boundary conditions. A uniform integration step of 2 fs was used during NPT simulations. Harmonic restrains in the tubulin backbone were utilized to avoid any structure distortion.

For the curved tubulin state targeted molecular dynamics was carried out to drive the tubulin conformation in the kip3-tubulin (straight) complex to a bent tubulin state using as a reference the conformation found in the 4HNA PDB (kinesin-1 – tubulin - DARPIN complex). The alpha carbons of residues 2-37, 49-437 in alpha tubulin, and 2-437 in beta tubulin were selected for the alignments. The TMD ramp was carried out during 5 ns and after that we run additional 25 ns of MD maintaining the restraints in the tubulin structure to avoid structure distortions. The integration step, thermostat and pressure control were the same than in the simulation with straight tubulin.

### Analysis of MD trajectories

The last 10 ns of equilibration simulations were used to perform trajectory analysis using VMD. The RMSD of equilibrated structures was measured and its convergence supports the stability of the conformers. Salt bridges were monitored if the donor-acceptor distance was less than 3.2 Å. Hydrogen bonds were counted if the donor-acceptor distance was less than 3.0 Å with a 20° cutoff for the angle formed by the donor, hydrogen, acceptor. Figures depicting molecular structures were created with UCSF Chimera (Pettersen et al., 2004) and VMD.

### Preparation of stabilized microtubules for functional assays

Tubulin was purified from bovine brains by two rounds of assembly and disassembly as described in (Castoldi and Popov, 2003). HiLyte 488, HiLyte 647, and biotin -labeled tubulin were purchased from Cytoskeleton. Motility assays were carried out using biotinylated, fluorescently labelled, Taxol-stabilized microtubules (10% labelled tubulin) as previously described (Su et al., 2013). Taxol was added to a concentration of 20 uM and was always present in assays with Taxol-stabilized microtubules.

To prepare GMPCPP-stabilized microtubules, microtubule seeds were first grown by incubating seed mix (20 uM tubulin, 0.2 uM biotin-tubulin, 5 uM fluorescein-tubulin, and 1 mM GMPCPP-Mg in BRB80 pH 6.9) at 37°C for 30 min. The elongation mix was made by diluting a seed mix aliquot at 1:10 ratio with BRB80 and 0.5 mM GMPCPP-MgCl. The seeds were added to the elongation mix at a 1:40 ratio. GMPCPP-stabilized microtubules were elongated for 3 h at 37°C before use.

### Single molecule assays for motility measurements

Flow chambers were prepared as previously described using double sided tape with a volume of ~10-15ul (Su et al., 2013). Chambers were incubated sequentially with the following solutions, interspersed with washes with BRB80 buffer + 20uM Taxol: 0.5mg/ml biotinylated-BSA (Vector Laboratories), 0.5mlg/ml streptavidin (Sigma), a 1:100 dilution of Taxol-stabilized microtubules in BRB80+Taxol. The last wash was performed with Reaction Buffer (BRB80 pH 7.2 with 0.5 mg/mL k-casein, 5% glycerol, 2 mM ATP-Mg, 20 uM Taxol and 100mM KCl, unless otherwise specified). Kip3 was diluted in reaction buffer at 4°C and introduced in the assay chamber in reaction buffer with an oxygen scavenging mix (0.2 mg/mL glucose oxidase, 0.035 mg/mL catalase, 25 mM glucose and 1mM DTT). For end-residence time experiments as a function of total motor concentration, unlabeled motor was diluted in reaction buffer and this was then mixed with TMR-labelled motor(<0.05 nM), introduced into the flow-cell and imaged. The TMR-labeled motors and HiLyte-488-labelled microtubules were imaged at room temperature in a Nikon Ti-E microscope with TIRF-illuminator, perfect-focus system with a 100x 1.49 NA Nikon oil immersion TIRF objective and 488 nm and 561 nm lasers (Agilent module). Images were recorded on an Andor DU-897 (EMCCD) camera at 100ms exposure and EMCCD gain of 150.

### TIRF assay to determine Kip3 lattice binding

Fluorescence intensity measurements to compare the relative binding of Kip3 constructs to Taxol-stabilized (GDP-like) microtubules were performed similarly to motility assays using TIRF imaging. Taxol-stabilized microtubules were grown with HiLyte-488 –labelled tubulin. Taxol-stabilized microtubules were introduced into the same flow chamber with the presence of 20uM Taxol in the reaction buffer. Taxol was added for microtubule stability as Kip3 is unable to depolymerize Taxol-stabilized microtubules. Kip3 was introduced at the desired concentrations and imaged as described in the motility assays. Analysis of background-subtracted Kip3 brightness on the microtubule lattices was performed in ImageJ. The Kip3 fluorescence intensity was calculated using all of the microtubules in a field of view. The ends (600 nm from the end) of stabilized or dynamic microtubules were excluded from the analysis to avoid end-specific effects.

### Depolymerization assays

Experimental conditions were similar to those for the motility assay, with the following modifications: GMPCPP-stabilized microtubules were used instead of Taxol-stabilized microtubules. Taxol was not included in the reaction mixtures. Wide-field images were acquired every 12 s for 12 min on an upright Zeiss AxioImager M1 microscope using a 63x objective and a Photometrics Cool Snap HQ camera (Su et al., 2013). Images were analyzed first by manually selecting microtubules using the same Labview software described in the motility assays. A custom Matlab software was used to automatically measure microtubule length in each frame. Depolymerization rates were calculated as changes in length over time.

For experiments that required normalization of motor binding to microtubules, fluorescence intensity of Kip3 motors bound to Taxol-stabilized microtubules was measured as described in the section: TIRF assay to determine Kip3 lattice binding. A binding curve of fluorescently labelled Kip3_448_ was first made by imaging the fluorescence intensity of Kip3_448_ at various input concentrations. A linear fit to the data points was performed in Matlab. This equation was then used for normalization of the protein concentration of motor variants needed to match the fluorescence intensity of Kip3_448_ on microtubules at 250nM input concentration. This value was then used to normalize the protein input in the depolymerization assay. For assays using ATP analogues, Kip3 variants were purified as previously described and exchanged to assay buffer in the corresponding ATP analogue and used at 1uM monomer as input.

### Tubulin Binding Affinity Measurements

Tubulin affinity measurements were performed in an Octet Red 384 instrument (ForteBio), all assays were performed in ATPase buffer without k-casein, with 10% sucrose, in ATP, unless otherwise stated. Monomeric Kip3 was immobilized on an anti-penta-HIS biosensor chip (HIS1K, ForteBio) at an input concentration of 100nM. Unbound motor was washed with ATPase buffer. Tubulin was diluted in ATPase buffer and introduced at a range of concentrations (10-fold the expected K_d_) of the motor construct. Upon reaching steady state, the sensor was automatically switched to a reaction buffer-only condition to measure tubulin off-rates. The data was analyzed using ForteBio software using both the measured k_on_ and k_off_ from a range of concentrations and steady state data to obtain the K_d_.

### Quantification and statistical analysis

Motor velocities and run lengths were analyzed using a software developed in Labview (National Instruments) for selecting microtubules, creating kymographs and tracking the position and brightness of motor tracks. Software was validated using manual analysis of kymographs in ImageJ. Kymographs of Kip3 runs and dwells were taken over the time course of the movie and were used to determine average rate of motility. Matlab was used to compile the data and perform distribution fittings for velocities (normal distribution), run lengths (exponential distribution) and plus-end dwell times (exponential distribution), using Scott’s rule for bin number, as previously described (Su et al., 2013). Statistical tests were performed using a two-tailed un-paired t-test.

## Author Contributions and Notes

H.A.S. and R.A.H.-L. contributed equally to this work and their authorship order was decided by a coin toss. H.A.S. performed the protein production and purification, as well as the biochemical and single-molecule assays. R.A.H.-L. performed the cryo-EM and the MD simulations and contributed to the single-molecule assays. E.S. contributed to the protein production and purification, as well as to the biochemical assays, R.Y.-R.W., performed the model building using Rosetta. H.A.S., R.A.H.-L., D.P. and A.E.L. analyzed the data. H.A.S., R.A.H.-L., D.P. and A.E.L. wrote and edited the manuscript.

The authors declare that there is no conflict of interest regarding the publication of this article.

## Acknowledgments

We thank N. Umbreit, B. W. Redwine and K. Toropova for critical reading of this manuscript. We are grateful to the NIC at HMS imaging center for the usage of TIRF microscopes. We acknowledge the center for molecular interactions at HMS-BCMP for the use of the Octet system. D. P. is supported by Howard Hughes Medical Institute and a National Institute of Health grant (GM61345). H.A.-S. was supported by an international fellowship from the Howard Hughes Medical Institute. R.H.-L. was supported by a Fundacion Mexico en Harvard Fellowship. We thank C. Sindelar (Yale), G. Alushin (The Rockefeller University), G. Lander (Scripps), R. Zhang (U Washington) and M. Cianfrocco (University of Michigan) for sharing processing scripts and helpful advice, M. Strauss (Harvard), Z. Yu (HHMI-Janelia) and J. de la Cruz (HHMI-Janelia) for help collecting EM data, T. Goddard and the Resource for Biocomputing, Visualization, and Informatics (UCSF) for helpful advice and for the use of the visualization resources. All the members of the Leschziner and Pellman Labs for advice and helpful discussions. EM data was collected at the EM facility of the Howard-Hughes Medical Institute at Janelia. Image processing and MD simulations were run on the Orchestra cluster at Harvard Medical School, the Odyssey cluster supported by the FAS Science Division Research Computing Group, Harvard University. This work also used the Extreme Science and Engineering Discovery Environment (XSEDE), which is supported by National Science Foundation grant number ACI-1053575 (AEL-RHL).

