## Supplementary Figures for "Multimodal tubulin binding by the yeast kinesin-8, Kip3, underlies its motility and depolymerization"

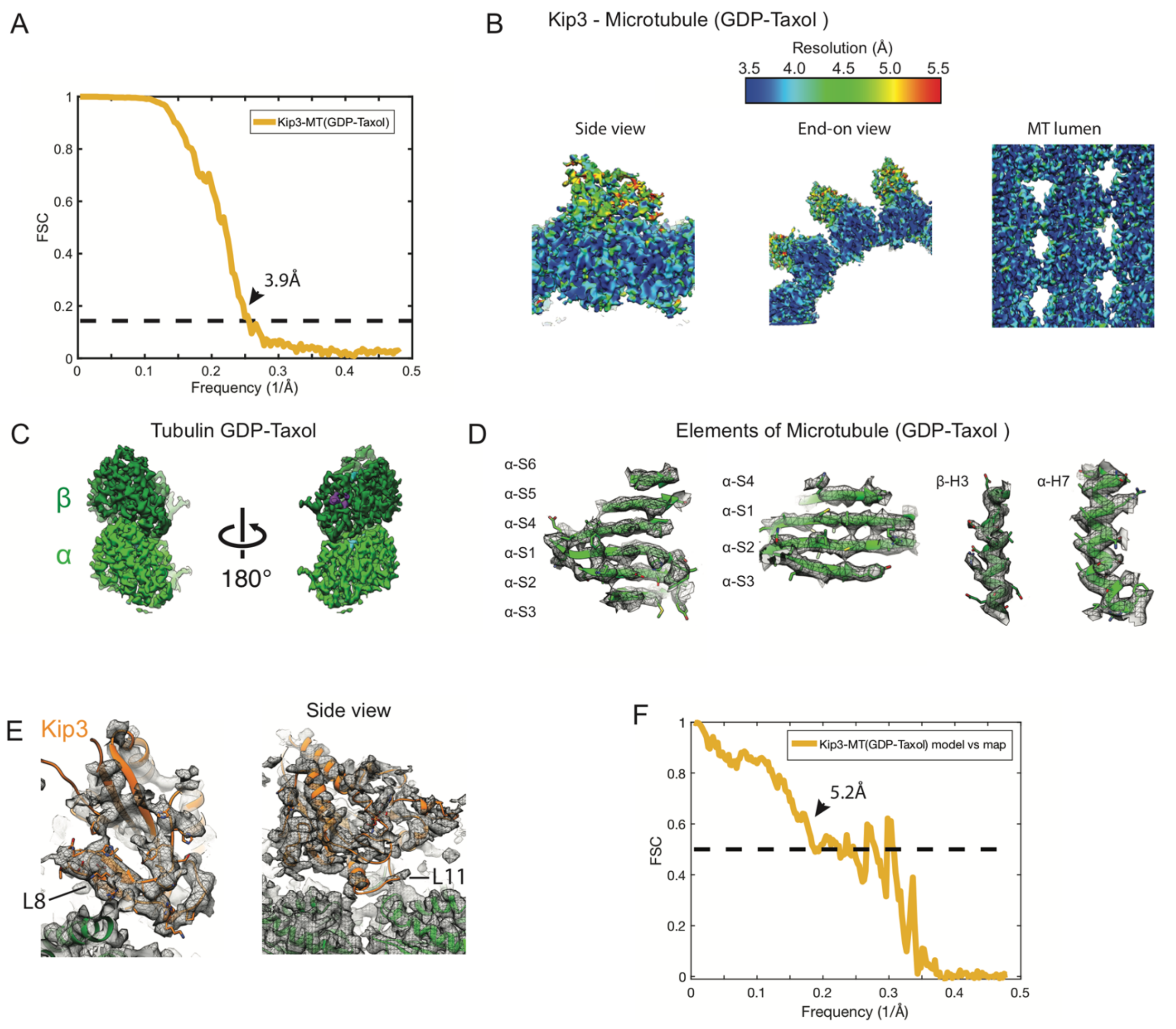


**Figure S1. Summary of Cryo-EM analysis for the Kip3-MT (GDP-Taxol) reconstruction.**

**(A)** Fourier Shell Correlation (FSC) curve for the Kip3-MT (GDP-Taxol) structure. **(B)** Local resolution of the cryo-EM map, as plotted by Bsoft (Heymann and Belnap, 2007). Side, end-on and MT lumen views are shown. The maps have been locally filtered to the indicated resolutions using Blocres (Cardone et al., 2013). The tubulin resolution ranges from 3.5Å in the core to 4 Å on the surface; the Kip3 resolution ranges from 4.5 Å to 5.5 Å. **(C)** Segmented densities corresponding to a tubulin dimer in the GDP-Taxol reconstruction. The map was sharpened and filtered according to the local resolution. **(D)** Representative densities of tubulin α-helices and β-sheets. **(E)** Close up of the region of the cryo-EM map and model corresponding to Kip3 (light orange). The densities shown are from the sharpened maps, filtered according to the local resolution. **(F)** Map-to-model FSC curve for Kip3-MT(GDP-Taxol).

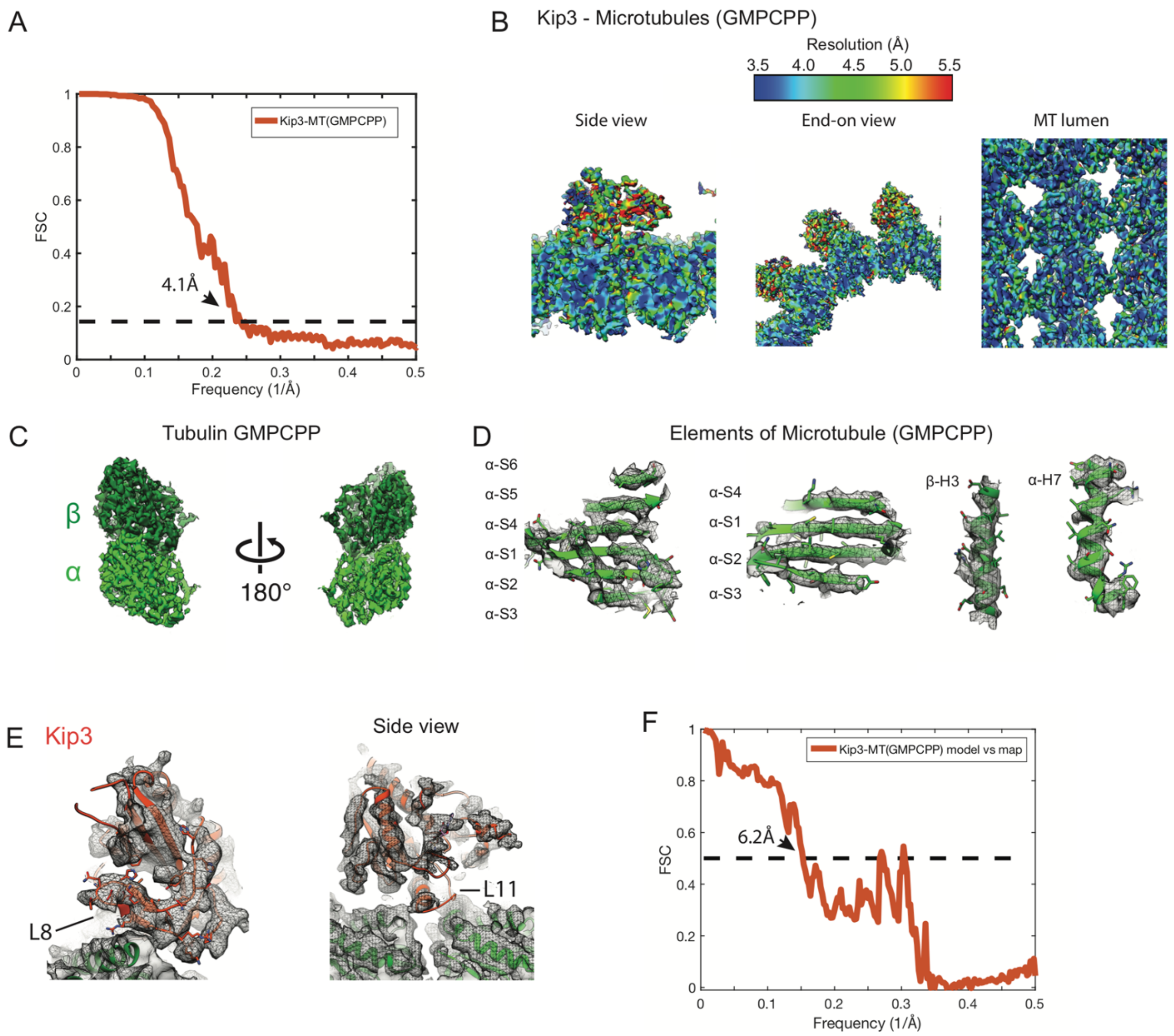


**Figure S2. Summary of Cryo-EM analysis for the Kip3-MT (GMPCPP) reconstruction.**

**(A)** Fourier Shell Correlation (FSC) curve for the Kip3-MT (GMPCPP) structure. **(B)** Local resolution of the cryo-EM map, as plotted by Bsoft (Heymann and Belnap, 2007). Side, end-on and MT lumen views are shown. The maps have been locally filtered to the indicated resolutions using Blocres (Cardone et al., 2013). The tubulin resolution ranges from 4.0 Å in the core to 4.5Å on the surface; the Kip3 resolution ranges from 4.5 Å to 6.0 Å. **(C)** Segmented densities corresponding to a tubulin dimer in the GMPCPP reconstruction. The map was sharpened and filtered according to the local resolution. **(D)** Representative densities of tubulin α-helices and β-sheets. **(E)** Close up of the region of the cryo-EM map and model corresponding to Kip3. The densities shown are from the sharpened maps, filtered according to the local resolution. **(F)** Map-to-model FSC curve for Kip3-MT(GMPCPP).

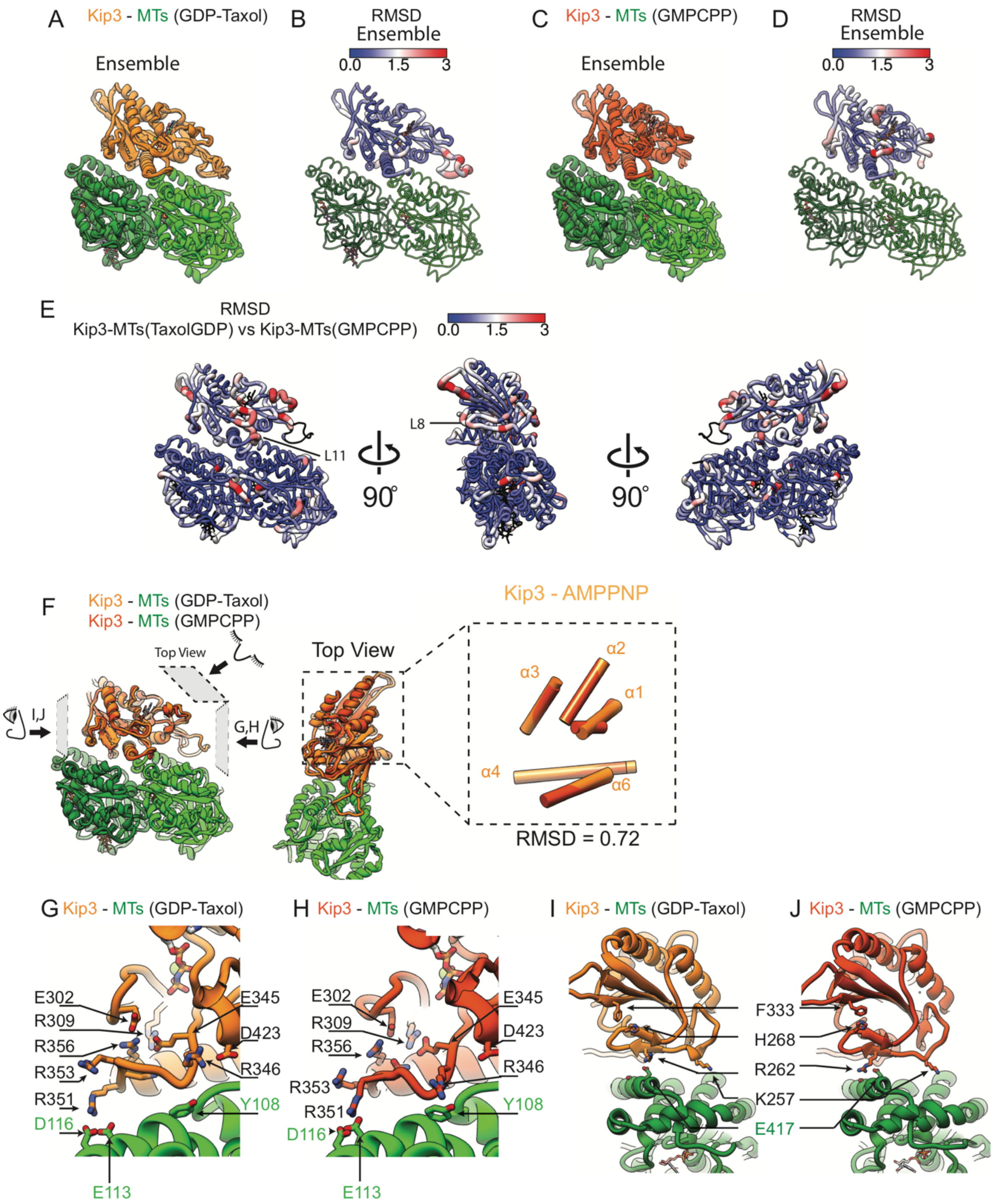


**Figure S3. Comparison of the Kip3-MT(GDP-Taxol) and Kip3-MT(GMPCPP) models.**
**(A)** Alignment of the top five Rosetta models for the Kip3-MT(GDP-Taxol) structure. **(B)** RMSD (Cα atoms) for the alignment in (A) plotted onto the structure. **(C)** Alignment of the top five Rosetta models for the Kip3-MT(GMPCPP) structure. **(D)** RMSD (Cα atoms) for the alignment in (C) plotted onto the structure. **(E)** Cα atom RMSD map between the top models for Kip3-MT(GDP-Taxol) and Kip3-MT(GMPCPP). L8 and L11, at the Kip3-tubulin interface show the largest RMSDs. **(F)** Comparison of the Kip3 structures on GDP-Taxol (Kip3 in light orange) and GMPCPP (Kip3 in dark orange). The atoms in the αβ tubulin backbone were used for alignment. Inset: Superposition of α-helices α1-α6 in the Kip3 structures shown from the top view and represented as cylinders. **(G, H)** Close-up views of interactions involving Kip3 L11 and tubulin in the GDP-Taxol and GMPCPP structures. **(I, J)** Close-up views of interactions involving Kip3 L8 and tubulin in the GDP-Taxol and GMPCPP structures.

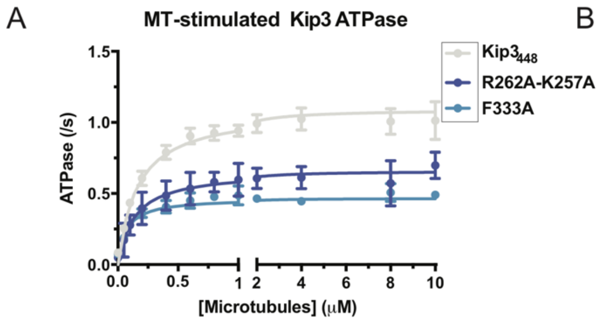


| | $k_{cat}(s^{-1})$ | $K_M(\mu M)$ |
|---|---|---|
| $Kip3_{448}$ | 1.09 +/- 0.36 | 0.16 +/- 0.36 |
| R262A-K257A | 0.66 +/- 0.04 | 0.13 +/- 0.04 |
| F333A | 0.47 +/- 0.01 | 0.07 +/- 0.02 |

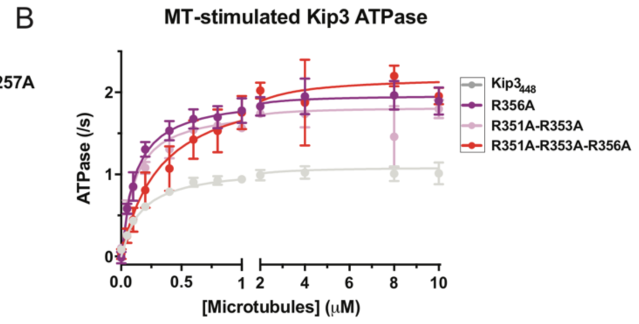


| | $k_{cat}(s^{-1})$ | $K_M(\mu M)$ |
|---|---|---|
| $Kip3_{448}$ | 1.09 +/- 0.36 | 0.16 +/- 0.36 |
| R356A | 1.97 +/- 0.02 | 0.11 +/- 0.01 |
| R351A-R353A | 1.82 +/- 0.03 | 0.11 +/- 0.01 |
| R351A-R353A R356A | 2.19 +/- 0.06 | 0.33 +/- 0.05 |

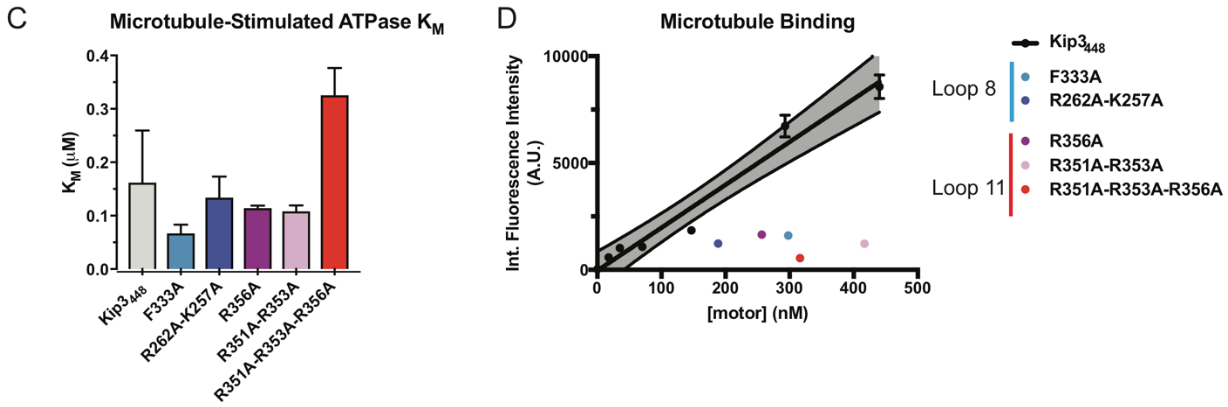


**Figure S4. Microtubule-stimulated ATPase activity, microtubule binding and representative kymographs of motility data of Kip3 WT and L8, L11 mutants.**

Microtubule-stimulated ATPase activity of $Kip3_{448}$ and $Kip3_{448}$ mutants in the presence of 1 mM ATP (mean +/- SEM). **(A)** Data for $Kip3_{488}$ and L8 mutants. **(B)** Data for $Kip3_{448}$ and L11 mutants. Tables show $K_{cat}$ and $K_M$ for $Kip3_{448}$ and mutants obtained from fitting the MT-stimulated ATPase curves to a Michaellis-Menten model. **(C)** Summary of $K_M$ for $Kip3_{448}$ and mutants. **(D)** Fluorescence intensity of $Kip3_{448}$ at different motor concentrations. Each point shows the fluorescent intensity of the Kip3 variant at the indicated concentration.

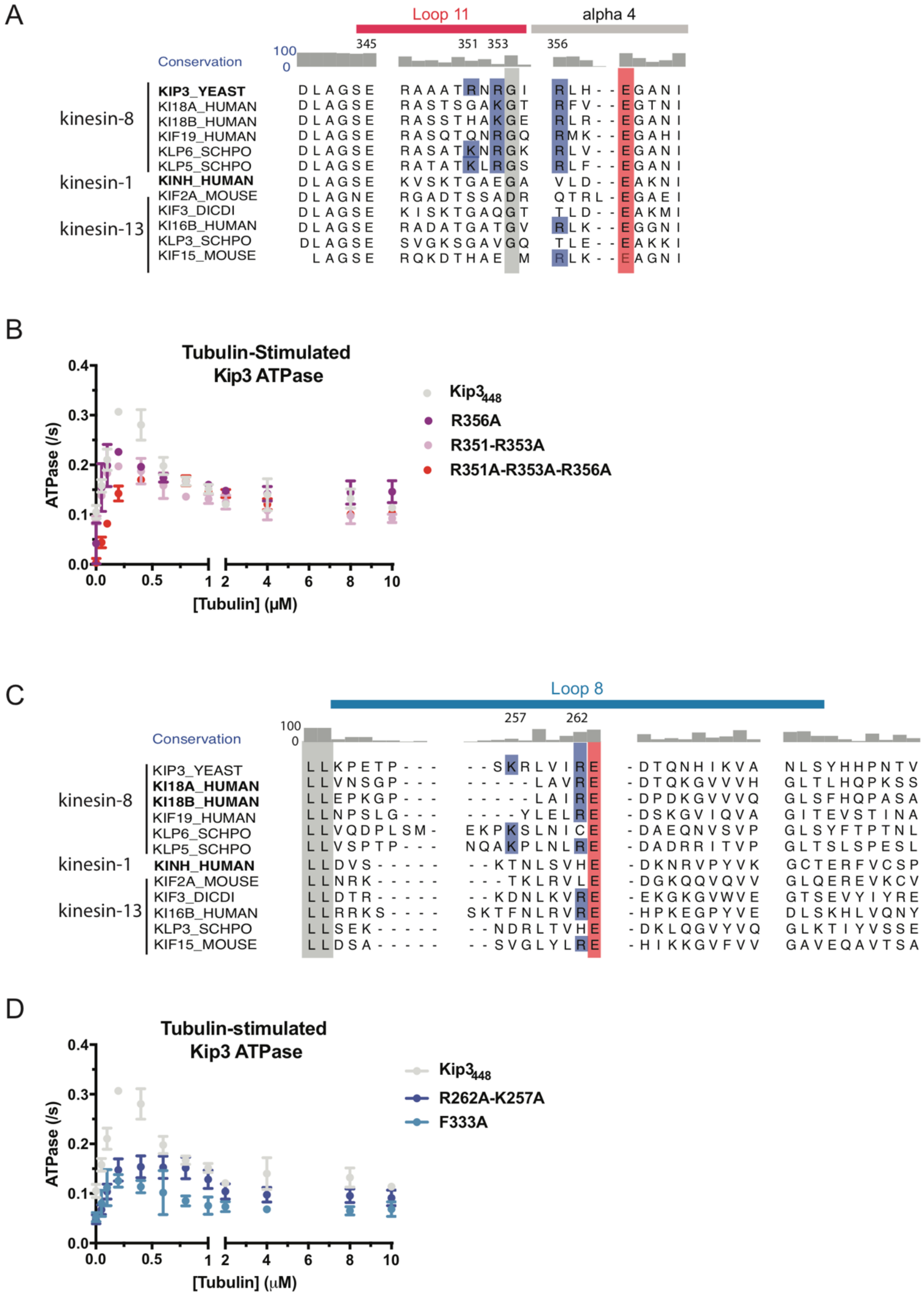


**Figure S5. Tubulin-stimulated ATPase activity data of Kip3 WT and mutants. Sequence alignment and conservation of L8 and L11 regions.**

Tubulin-stimulated ATPase activity of $Kip3_{488}$, $Kip3_{488}$ -L8 mutants **(A)** and $Kip3_{488}$-L11 mutants **(B)** in the presence of 1 mM ATP (mean +/- SEM). Sequence alignment and conservation of Kip3 residues in the L8 region **(C)** and L11 and α4 regions **(D)**.

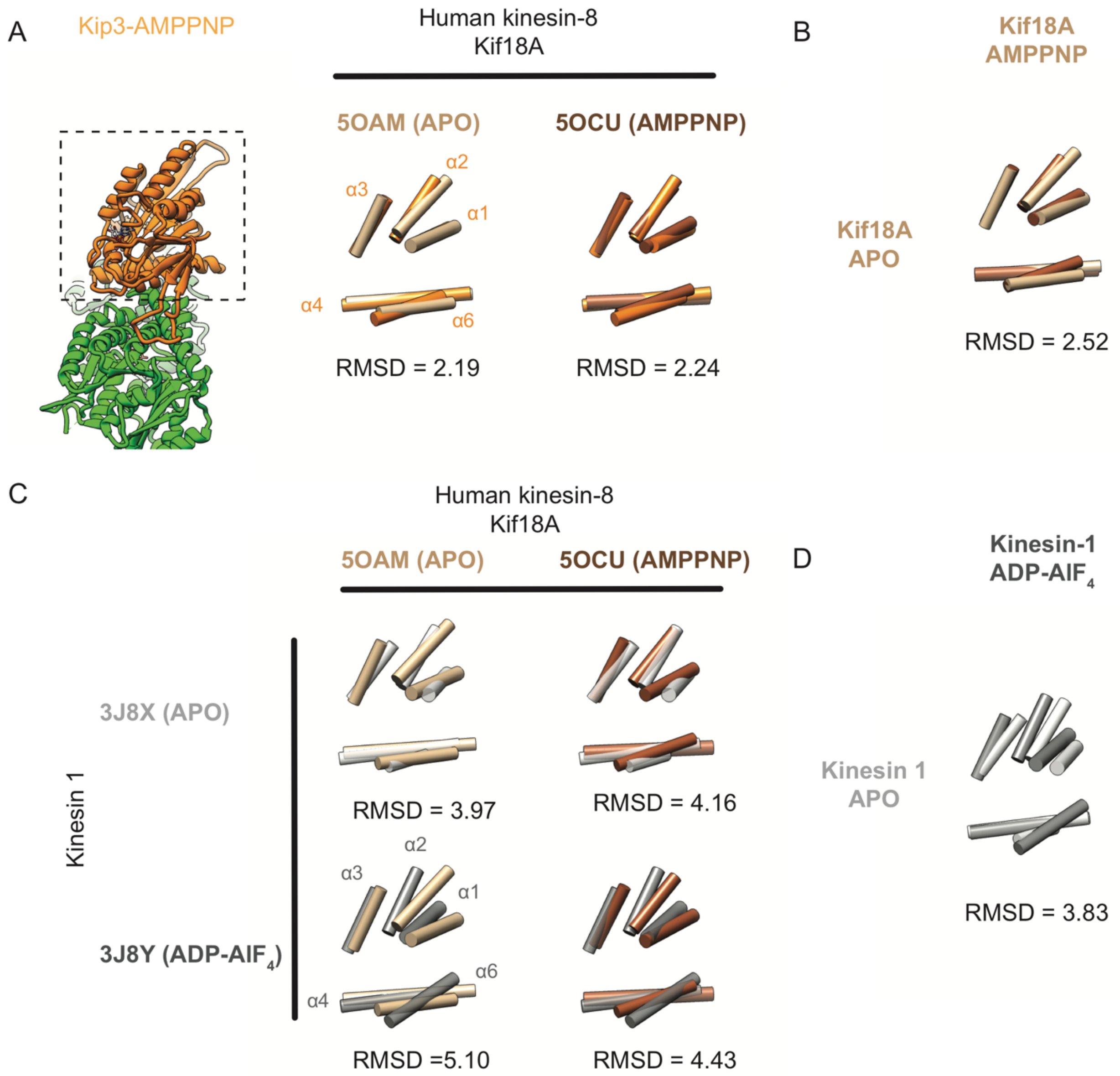


**Figure S6. Comparison of the Kif18A in the APO and AMPPNP states with Kip3(AMPPNP) and Kinesin-1 in the APO and ADP-AlF$_4$.**

The APO-like AMPPNP-bound state is conserved in the human kinesin-8 structures. **(A)** Front view of the Kip3(AMPPNP) structure. Comparison of the Kip3(AMPPNP) (light orange) with human kinesin-8 (Kif18A) in the APO (light brown, PDB 5OAM) or AMPPNP (dark brown, PDB 5OCU) states. Superposition of alpha-helices α1-α6 in the Kip3 structures shown from the top view and represented as cylinders. The atoms in the αβ-tubulin backbone were used for alignment. **(B)** Comparison of the Kif18A structures in the APO (light brown) and AMPPNP (dark brown) states. **(C)** Comparison of the human kinesin-8 (Kif18A) in the APO (light brown, PDB 5OAM) or AMPPNP (dark brown, PDB 5OCU) states with kinesin-1 APO (white, PDB 3J8X) or kinesin-ATP-like (gray, PDB 3J8Y). **(D)** Comparison of the kinesin-1 structures in the APO (light gray) and ADP-AlF$_4$ (dark gray) conformations.

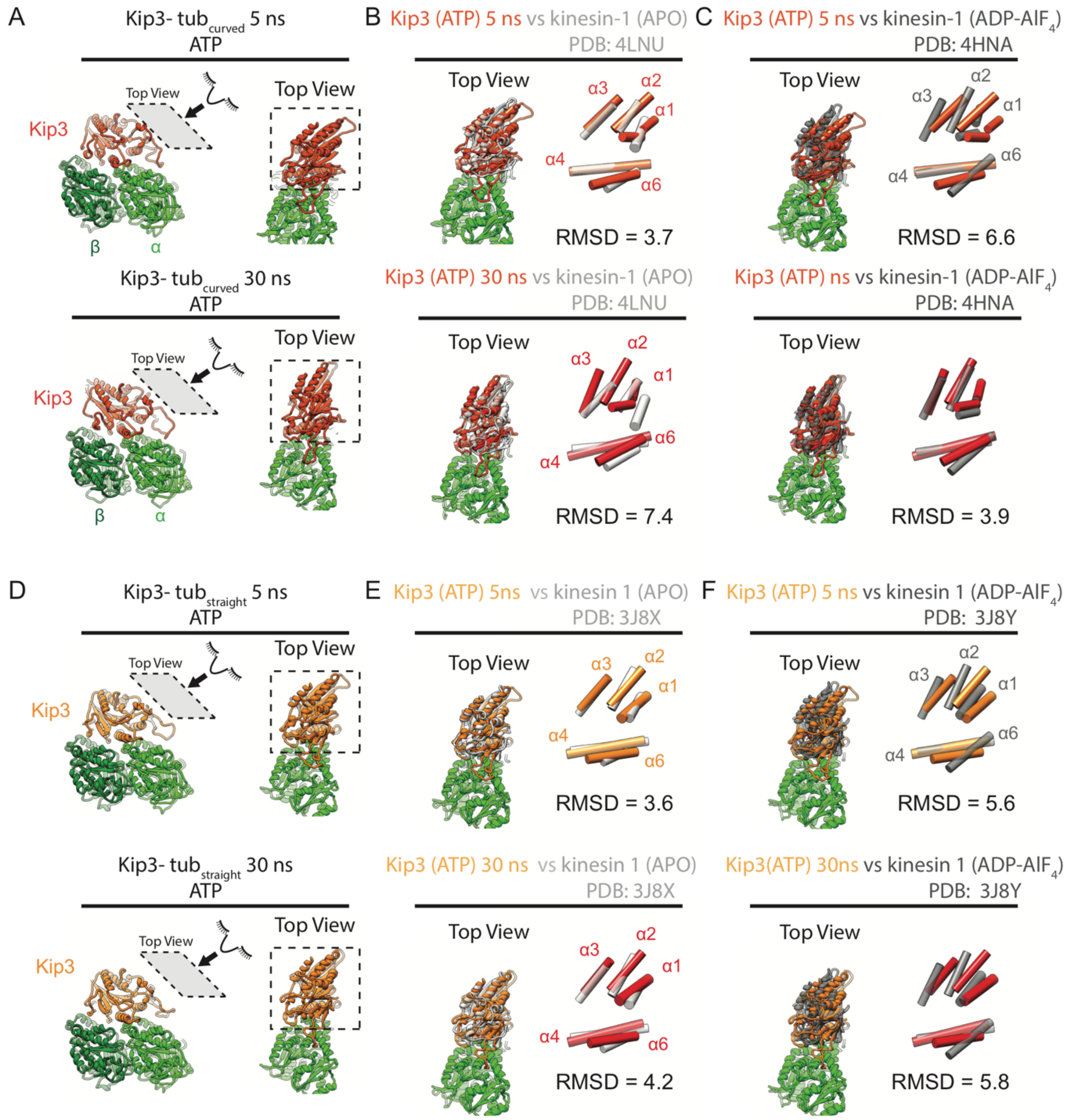


**Figure S7. Structural analysis of Kip3 conformations when bound to different tubulin conformations during MD trajectories.**

Molecular dynamics simulations suggest that binding to curved tubulin stabilizes the ATP-like conformation of Kip3. **(A)** Snapshots of Kip3 (ATP) bound to tubulin during targeted molecular dynamics simulations (transition from straight to curved tubulin) at 5 ns (top) and 30 ns (bottom). Comparison of top views of Kip3 at 5ns (top) and 30 ns (bottom) with kinesin-1 in the **(B)** APO (light gray, PDB 4LNU) or **(C)** ADP-$AlF_4$ (dark gray, PDB 4HNA) states, in both structures kinesin-1 is bound to curved tubulin. The core alpha helices (α1-α6) of Kip3 and kinesin-1 are shown as cylinders. **(D-F)** Binding to straight tubulin is not sufficient to stabilize the ATP-like state of Kip3. Control experiments (straight tubulin -> straight tubulin) show that the conformational switch to the ATP-like state of Kip3 does not take place in the absence of curved tubulin **(D)** Snapshots of Kip3 (ATP) bound to straight tubulin (MT) during molecular dynamics simulation (fixed straight tubulin) at 5ns (top) and 30 ns (bottom). Comparison of top views of Kip3 conformations at 5ns (top) and 30 ns (bottom) with kinesin-1 in the **(E)** APO (light gray, PDB 3J8X) or **(F)** ADP-$AlF_4$ (dark gray, PDB 3J8Y) states, kinesin-1 is bound to straight tubulin (MTs) in both structures. The core alpha helices (α1-α6) of Kip3 and kinesin-1 are shown as cylinders.

A

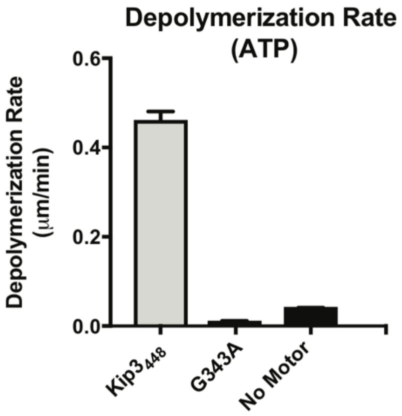


**Figure S8. Depolymerization rates of GMPCPP-stabilized MTs of Kip3 and G343A mutant.**
**(A)** Depolymerization rates of GMPCPP-stabilized MTs with 1μM of the indicated constructs in the presence of 1mM ATP, except for $Kip3_{438}$ (grey), which was used at 250 nM due to its high depolymerization rate (mean +/- sem).
