## Supplementary Tables for "Multimodal tubulin binding by the yeast kinesin-8, Kip3, underlies its motility and depolymerization"

**Table S1. Data collection, refinement and model statistics**

| | Kip3-MTs (Taxol)<br>EMD-24666<br>PDB: 7RS5 | Kip3-MTs (GMPCPP)<br>EMD-24667<br>PDB: 7RS6 |
|---|---|---|
| **Data Collection** | | |
| Microscope | Titan Krios | FEI Polara |
| Voltage (kV) | 300 | 300 |
| Detector | K2-summit | K2-summit |
| Frames | 20 | 20 |
| Exposure time (s) | 4 | 5 |
| Total dose ($e^{-1}/Å^2$) | 40 | 40 |
| Processed Micrographs | 1194 | 643 |
| Pixel size of Processed data | 1.04 | 0.98 |
| Number of extracted particles | 67040 | 29090 |
| 14 PF Particles | 23955 | 22264 |
| Final 14 PF particles | 14934 | 21788 |
| Asymmetric units in 14 PF | 209,076 | 305032 |
| Resolution 14PF map | 3.9 | 4.1 |
| Twist per unit (degrees) | -25.76 | -25.76 |
| Rise per unit (Å) | 8.55 | 8.90 |
| **Refinement** | | |
| Resolution (Å) | 3.9 | 4.1 |
| Map sharpening B-factor ($Å^2$) | 100 | 150 |
| Overall correlation of the residues to the map (RSCC) | 0.683 | 0.764 |
| **RMS deviation** | | |
| Bonds (Å) | 0.0115 | 0.0111 |
| Angles (°) | 1.17 | 1.15 |
| **Model Geometry** | | |
| Molprobity score | 1.38 | 1.44 |
| Clashscore (all atoms) | 1.75 | 2.02 |
| Favored rotamers (%) | 98.26 | 98.14 |
| **Ramachandran plot** | | |
| Favored (%) | 92.95 | 92.33 |
| Outliers (%) | 2.10 | 1.29 |

**Data Availability**

The cryo-EM maps for Kip3 (AMPPNP) bound to either Taxol-stabilized microtubules or GMPCPP-stabilized microtubules are deposited in the EMDB as EMD-24666 and EMD-24667, respectively. The corresponding molecular models are deposited in the PDB as 7RS5 and 7RS6

**Table S2. Prominent Interactions in Kip3-Microtubule models**

| Donor | Acceptor | Distance (Å) MT-Taxol Model | Distance (Å) MT-GMPCPP Model |
|---|---|---|---|
| R262 Kip3 | E420 β-Tubulin | 5.21 | **2.82** |
| R262 Kip3 | S423 β-Tubulin | 8.66 | **3.29** |
| R309 Kip3 | E345 Kip3 | 4.89 | **2.75** |
| R346 Kip3 | D423 Kip3 | **3.13** | 7.92 |
| R346 Kip3 | Y108 α-Tubulin | 6.10 | **2.98** |
| R351 Kip3 | E113 α-Tubulin | **2.91** | 5.25 |
| R351 Kip3 | D116 α-Tubulin | 8.28 | **2.83** |
| N352 Kip3 | E345 Kip3 | 3.80 | **2.88** |
| R353 Kip3 | E113 α-Tubulin | 4.25 | 7.28 |
| R356 Kip3 | E302 Kip3 | **2.76** | **2.80** |
| R356 Kip3 | E345 Kip3 | **2.86** | 6.37 |
| H268 Kip3 | F333 Kip3 | 3.72 | 3.93 |

Numbers in pink denote hydrogen bonds as found by geometrical and distance search using UCSF Chimera.

**Table S3. Intermolecular Interactions in Kip3 - tubulin models during molecular dynamics simulations (30 ns)**

| | | **Straight tubulin (MT)** | **Curved tubulin (both)** | **Curved tubulin (alpha)** | **Curved tubulin (beta)** |
|---|---|---|---|---|---|
| Unique salt bridges in 30 ns | | 181 | 206 | 193 | 194 |
| Unique hbonds in 30 ns | | 83 | 84 | 91 | 86 |
| Unique hbonds above 10% occupancy (3 ns) | | 18 | 23 | 22 | 19 |
| **Donor - Acceptor** | **Kip3 Domain** | **Straight tubulin (MT) % occupancy** | **Curved tubulin (both) % occupancy** | **Curved tubulin (alpha) % occupancy** | **Curved tubulin (beta) % occupancy** |
| K257 Kip3 - E420 β-Tubulin | Loop 8 | - | 23.27 | - | - |
| K257 Kip3 - E417 β-Tubulin | Loop 8 | - | 18.9 | - | 24.93 |
| K257 Kip3 - E159 β-Tubulin | Loop 8 | 12.27 | 8.00 | 36.30 | - |
| R258 Kip3 - D414 β-Tubulin | Loop 8 | 5.87 | 19.5 | 2.97 | 14.37 |
| R258 Kip3 - E417 β-Tubulin | Loop8 | 87.2 | 7.5 | 76.03 | 1.47 |
| R262 Kip3 - E414 β-Tubulin | Loop 8 | 1.30 | - | 21.17 | 1.47 |
| R262 Kip3 - E417 β-Tubulin | Loop 8 | - | 69.7 | - | |
| R262 Kip3 - E420 β-Tubulin | Loop 8 | 0.43 | 32.47 | 2.53 | 4.33 |
| D264 Kip3 - T419 β-Tubulin | Loop 8 | - | 28.97 | 1.93 | - |
| D264 Kip3 - S423 β-Tubulin | Loop 8 | - | - | 9.00 | 41.13 |
| S344 Kip3 - E414 α-Tubulin | **Loop 11** | 0.07 | 60.2 | 53.30 | 8.50 |
| R346 Kip3 - E414 α-Tubulin | **Loop 11** | 74.63 | 58.63 | - | 3.27 |
| R346 Kip3 - E417 α-Tubulin | **Loop 11** | - | 13.7 | - | 13.7 |
| R351 Kip3 - D116 α-Tubulin | **Loop 11** | 4.17 | 42.57 | 4.43 | 41.53 |
| R351 Kip3 - E113 α-Tubulin | **Loop 11** | - | - | - | 17.50 |
| R351 Kip3 - E155 α-Tubulin | **Loop 11** | 10.1 | - | - | - |
| Y108 α-Tubulin - N352 Kip3 | **Loop 11** | - | 29.8 | - | 4.13 |

| | | | | | |
|---|---|---|---|---|---|
| R353 Kip3 - E113 α-Tubulin | **Loop 11** | 6.03 | 2.13 | 47.67 | 61.43 |
| R353 Kip3 - D116 α-Tubulin | **Loop 11** | 7.43 | - | - | - |
| K139 Kip3 - D199 α-Tubulin | **Loop 2** | - | 41.6 | - | - |
| R144 Kip3 - E196 α-Tubulin | **Loop 2** | 70.33 | 73.03 | 82.67 | 85.93 |
| R145 Kip3 - E420 α-Tubulin | **Loop 2** | 65.43 | 67.83 | 42.17 | 54.00 |
| R145 Kip3 - E423 α-Tubulin | **Loop 2** | 37.27 | 31.17 | 14.10 | 42.30 |
| R145 Kip3 - D424 α-Tubulin | **Loop 2** | 87.43 | 78.17 | 85.23 | 53.10 |
| R146 Kip3 - E420 α-Tubulin | **Loop 2** | 21.87 | 27.33 | 42.80 | 65.97 |
| R245 Kip3 - E159 β-Tubulin | Beta 5 | 30.43 | 48.00 | 38.07 | 45.77 |
| H387 kip3 - Q434 β-Tubulin | Alpha 5 | 12.17 | 0.73 | 23.97 | 7.60 |
| R391 Kip3 - E431 β-Tubulin | Alpha 5 | 64.33 | 82.23 | 91.63 | 59.50 |
| K394 Kip3 - E196 β-Tubulin | Alpha 5 | 19.37 | 12.63 | 13.80 | 18.57 |
| R397 Kip3 - E196 β-Tubulin | Alpha 5 | 36.73 | 27.77 | 68.03 | 82.33 |
| R397 Kip3 - E420 β-Tubulin | Alpha 5 | 39.57 | 24.37 | 3.50 | - |
| R384 Kip3 - D437 β-Tubulin | Loop 12 | 31.13 | - | - | - |
| K430 Kip3 - E420 α-Tubulin | Alpha 6 | 15.87 | 2.37 | 16.43 | 1.33 |
| R434 Kip3 - E415 α-Tubulin | Alpha 6 | 41.9 | 13.5 | 67.57 | 18.33 |

**Table S4. Intramolecular Interactions in Kip3 - tubulin models during the last 10 ns of molecular dynamics simulations**

| Kip3 – Kip3 | Location | Straight tubulin (MT) % occupancy | Curved tubulin (both) % occupancy | Curved tubulin (alpha) % occupancy | Curved tubulin (beta) % occupancy |
|---|---|---|---|---|---|
| R309 - E345 | Catalytic residues | 40.56 | 91.11 | 86.81 | 86.61 |
| R309 - E359 | Catalytic residues | 44.76 | - | - | - |
| S344 - E424 | Loop 11 - alpha 6 | 55.84 | - | - | - |
| R346 - E424 | Loop 11 - alpha 6 | 84.02 | - | 78.22 | 82.72 |
| R346 - D423 | Loop 11 - alpha 6 | - | 1.70 | 12.09 | 7.89 |
| R356 - E345 | Loop 11 | 72.73 | 93.21 | 10.49 | - |
| R356 - E359 | Loop 11 | - | 64.94 | 17.58 | 86.01 |
| N352 - A347 | Loop 11 | 40.26 | - | - | - |
| N352 - A348 | Loop 11 | 8.79 | - | 14.59 | - |
| ~ | Loop 11 | 68.53 | - | - | - |
| R356 - E302 | Loop 11 | 31.37 | - | - | - |
| R365 - E244 | Alpha 4 | - | 78.92 | 0.4 | 61.74 |
| R146 - D423 | Loop 2 - alpha 6 | 65.53 | 83.82 | 12.39 | 7.29 |

**Table S5. Intermolecular Interactions in Kip3 - tubulin models during molecular dynamics simulations (last 10 ns)**

| | | Straight Tubulin | Curved tubulin (both) | Curved tubulin (alpha) | Curved tubulin (beta) |
|---|---|---|---|---|---|
| Unique hbonds in 10 ns | | 41 | 47 | 53 | 46 |
| Unique hbonds above 10% occupancy (1 ns) | | 17 | 20 | 22 | 16 |
| **Donor - Acceptor** | **Kip3 Domain** | **Straight tubulin (MT) % occupancy** | **Curved tubulin (both) % occupancy** | **Curved tubulin (alpha) % occupancy** | **Curved tubulin (beta) % occupancy** |
| K257 Kip3 - E420 β-Tubulin | Loop 8 | - | 58.94 | - | - |
| K257 Kip3 - E417 β-Tubulin | Loop 8 | - | 48.65 | - | 22.08 |
| K257 Kip3 – E159 β-Tubulin | Loop 8 | - | - | 37.76 | - |
| R258 Kip3 - E417 β-Tubulin | Loop8 | 86.61 | 11.69 | 54.95 | - |
| R258 Kip3 - D414 β-Tubulin | Loop 8 | 13.59 | 31.07 | 3.40 | - |
| R262 Kip3 - E414 β-Tubulin | Loop 8 | - | - | 9.19 | - |
| R262 Kip3 - E417 β-Tubulin | Loop 8 | - | 69.83 | - | - |
| R262 Kip3 - E420 β-Tubulin | Loop 8 | - | 26.47 | 2.30 | - |
| D264 Kip3 - T419 β-Tubulin | Loop 8 | - | 56.64 | - | - |
| D264 Kip3 - S423 β-Tubulin | Loop 8 | | | 5.09 | 47.45 |
| S344 Kip3 - E414 α-Tubulin | **Loop 11** | - | 58.34 | 56.44 | - |
| R346 Kip3 - E414 α-Tubulin | **Loop 11** | 86.51 | 81.72 | - | 9.79 |
| R346 Kip3 - E417 α-Tubulin | **Loop 11** | - | 11.89 | - | - |
| R351 Kip3 - D116 α-Tubulin | **Loop 11** | 12.49 | 79.12 | 12.39 | - |
| R351 Kip3 - E113 α-Tubulin | **Loop 11** | | | | 52.45 |
| R351 Kip3 – E155 α-Tubulin | **Loop 11** | 4.40 | - | - | - |
| R353 Kip3 - E113 α-Tubulin | **Loop 11** | - | - | 29.27 | 22.48 |

| | | | | | |
|---|---|---|---|---|---|
| K139 Kip3 - D199 α-Tubulin | **Loop 2** | - | 44.56 | - | - |
| R144 Kip3 - E196 α-Tubulin | **Loop 2** | 84.02 | 59.44 | 87.01 | 89.01 |
| R145 Kip3 - E420 α-Tubulin | **Loop 2** | 66.33 | 76.72 | 43.56 | 46.75 |
| R145 Kip3 - E423 α-Tubulin | **Loop 2** | 43.66 | 44.46 | 19.98 | 25.27 |
| R145 Kip3 - D424 α-Tubulin | **Loop 2** | 91.9 | 53.35 | 81.82 | 68.83 |
| R146 Kip3 - E420 α-Tubulin | **Loop 2** | | | 52.05 | 67.43 |
| H387 kip3 - Q434 β-Tubulin | Alpha 5 | 11.49 | - | 23.28 | 0.6 |
| R391 Kip3 - E431 β-Tubulin | Alpha 5 | 53.65 | 82.62 | 93.21 | 69.83 |
| K394 Kip3 - E196 β-Tubulin | Alpha 5 | 32.07 | - | 9.89 | 16.78 |
| R397 Kip3 - E196 β-Tubulin | Alpha 5 | 24.08 | 48.15 | 84.02 | 83.42 |
| R397 Kip3 - E420 β-Tubulin | Alpha 5 | 43.46 | 18.18 | - | - |
| R245 Kip3 - E159 β-Tubulin | Beta 5 | 16.88 | 45.95 | 51.35 | 44.66 |
| K430 Kip3 – E420 α-Tubulin | Alpha 6 | 26.07 | - | 12.29 | - |
| R434 kip3 – E415 α-Tubulin | Alpha 6 | 40.66 | - | 68.33 | 27.57 |
| R384 Kip3 - D437 β-Tubulin | Loop 12 | 36.76 | - | - | - |
